# Genetic Optimisation of Bacteria-Induced Calcite Precipitation in *Bacillus subtilis*

**DOI:** 10.1101/2021.08.17.456648

**Authors:** Timothy Dennis Hoffmann, Kevin Paine, Susanne Gebhard

## Abstract

**Background:** Microbially induced calcite precipitation (MICP) is an ancient property of bacteria, which has recently gained considerable attention for biotechnological applications. It occurs as a by-product of bacterial metabolism and involves a combination of chemical changes in the extracellular environment, e.g. pH increase, and presence of nucleation sites on the cell surface or extracellular substances produced by the bacteria. However, the molecular mechanisms underpinning MICP and the interplay between the contributing factors remain poorly understood, thus placing barriers to the full biotechnological and synthetic biology exploitation of bacterial biomineralisation.

**Results:** In this study, we adopted a bottom-up approach of systematically engineering *Bacillus subtilis*, which has no detectable intrinsic MICP activity, for biomineralisation. We showed that heterologous production of urease can induce MICP by local increases in extracellular pH, and this can be enhanced by co-expression of urease accessory genes for urea and nickel uptake, depending on environmental conditions. MICP can be strongly enhanced by biofilm-promoting conditions, which appeared to be mainly driven by production of exopolysaccharide, while the protein component of the biofilm matrix was dispensable. Attempts to modulate the cell surface charge of *B. subtilis* had surprisingly minor effects, and our results suggest this organism may intrinsically have a very negative cell surface, potentially predisposing it for MICP activity.

**Conclusions:** Our findings give insights into the molecular mechanisms driving MICP in an application-relevant chassis organism and the genetic elements that can be used to engineer *de novo* or enhanced biomineralisation. This study also highlights mutual influences between the genetic drivers and the chemical composition of the surrounding environment in determining the speed, spatial distribution and resulting mineral crystals of MICP. Taken together, these data pave the way for future rational design of synthetic precipitator strains optimised for specific applications.

## Background

Microorganisms contribute to the formation of Earth’s landscapes through mineral deposits via a process known as microbially induced calcite precipitation (MICP). Over the past 30 years, bacteria capable of MICP have been at the basis of innovative biotechnologies arising within civil engineering sectors. Example applications include consolidation of soils, bioremediation of heavy metal contamination, restoration of degrading stone surfaces, self-healing concrete and carbon dioxide sequestration (1–3). More recently, MICP has contributed to the emergence of Engineered Living Materials as a new discipline at the interface between materials science and synthetic biology. Here, the targeted formation of minerals is envisaged to contribute to the *de novo* formation or assembly of engineered materials with living cells as their basis (4). To enable tailoring of material formation and properties via the rational design process at the heart of synthetic biology and to fully exploit the potential of bacterial biomineralisation for materials science and biotechnology, a comprehensive understanding of the genetic drivers of MICP is essential.

MICP occurs as a by-product of bacterial metabolism, which creates a microenvironment that favours the precipitation of calcium cations and carbonate anions in the form of mineral calcium carbonate, predominantly calcite. This process is dependent on changes in pH, ion concentrations and availability of cell surface nucleation sites (5). The best-understood bacterial process driving MICP is ureolysis, which accounts for the fastest route of precipitation via a rapid increase in pH due to the release of ammonia (6, 7). Additionally, cellular aspects of the bacteria are important, with cell surface charge serving as a mechanism for attracting Ca^2+^ ions and providing nucleation sites to initiate crystal formation (5, 8). In a similar context, extracellular substances produced by bacteria can contribute to modulation of the microenvironment and provision of conditions favourable for precipitation. Of these, biofilm formation is the most relevant process for application (9, 10). However, it is not clear how much each of these elements contributes to MICP, and even less so how the individual contributors interact to determine the overall precipitation behaviour of a bacterium. To optimise and control MICP via genetic engineering, such knowledge is required.

Most microbial urease enzymes are nickel metalloenzymes, characteristically composed of three structural subunits encoded by the genes *ureA*, *ureB* and *ureC* (11). For synthesis of the catalytically active urease, additional accessory proteins are required that ensure correct folding and assembly, typically encoded by *ureD*, *ureF*, *ureG* and *ureE* (11). In addition, less essential accessory genes may be present that encode nickel permeases (e.g. *ureH* or *ureJ*) or urea transporters (12–15). The latter can be the high-affinity UrtABCDE system or the proton-gated UreI permease (14, 16), while some bacteria harbour genes for the eukaryotic-type low-affinity transporter UT. This breadth of accessory functions gives rise to a diversity of urease gene clusters with presumably a range of impacts on ureolytic ability and performance.

Recent studies that explored heterologous urease expression to engineer MICP have moved the urease genes of the strongly ureolytic and MICP-capable bacterium *Sporosarcina pasteurii* into *Escherichia coli* (17–19) or *Pseudomonas aeruginosa* (20). While this provided valuable proof-of-concept insights, these host organisms did not reflect the final target for biotechnological application. Of more direct relevance to application are non-pathogenic, spore-forming Gram-positive species, as they are both safer and able to survive for a long time as spores. Additionally, questions remain on the minimum set of genes of the urease cluster required to maximise precipitation. Therefore, our aim was to use the Gram-positive model and industrially relevant bacterium *Bacillus subtilis* as a chassis to determine the components of the urease gene cluster required to induce calcite precipitation. Its low intrinsic urease and MICP activity was expected to provide an ideal background to detect small changes in activity and provide a high-resolution understanding of the genes and traits that contribute to MICP.

Apart from metabolism as a driver for mineral precipitation, bacterial surfaces also play a role. Surface charge is governed by functional groups such as carboxyl and phosphate groups. In Gram-positive bacteria, a major contributor to surface charge are the teichoic acids, which are rich in phosphate groups in their backbone and thus impart an overall negative charge (21). Bacteria can actively control this charge through D-alanylation of teichoic acids, catalysed by the DltABCD system, which renders the cell surface less negatively charged and, for example, contributes to the bacterium’s defence against cationic antimicrobial compounds (21–23). The existence of a discrete genetic element to control surface charge, namely the *dltABCD* operon, provides a potential means of controlling MICP via surface charge modification through manipulation of the Dlt system. However, it was not known to what degree this system impacts on surface charge in *B. subtilis* and whether this could be exploited to influence precipitation ability.

Beyond the immediate cell surface, biofilms create an extracellular microenvironment that can be beneficial to precipitation through trapping of ions as well as providing favourable functional groups for crystal nucleation (9, 10). *B. subtilis* biofilms contain extracellular polymeric substances (EPS) predominantly comprised of polysaccharides and proteins (24, 25). Polysaccharide production is facilitated by the 15-gene *epsA-O* operon, while production of proteinaceous fibres is encoded by *tasA* of the *tapA-sipW-tasA* operon (26, 27). The contribution of biofilm formation to biomineralisation has been investigated previously, although the role of individual biofilm matrix components in MICP is not well understood. Past work studying the interactions between biomineralisation and biofilm formation in *B. subtilis* NCIB3610 showed that both *epsH* and *tasA* were required for wild-type localisation of calcite precipitates within the bacterial colony (28). However, the deletion strains used in this study showed a markedly reduced complexity of biofilm architecture, and it was therefore not clear whether the loss of biofilm structure or the loss of specific EPS components was responsible for impaired calcite localisation and to what degree.

To provide a more detailed understanding of the genetic drivers of MICP and how their interactions govern biomineralisation in an application-relevant chassis organism, we here investigated how urease activity, extracellular biofilm composition and cell surface charge can be used to engineer *Bacillus subtilis* for MICP activity. Using genetic manipulation coupled to functional strain characterisation, qualitative and quantitative assessment of MICP and imaging of precipitates, we show that heterologous expression of the ureolytic pathway, particularly in conjunction with urease accessory genes, and modulation of biofilm production, but not of Dlt activity, facilitated MICP in a formerly non-precipitating host. The outcomes of our study provide mechanistic insights to guide rational genetic engineering approaches for MICP. This may lead not only to optimised precipitator strains for current applications, but to development of the careful control over the MICP process required to design future engineered living materials.

## Results

### *Bacillus subtilis* W168 can be engineered to display strong ureolytic activity

The first step in developing synthetic MICP activity in *B. subtilis* W168 was to enhance its ureolytic activity. *B. subtilis* possesses the core structural genes for urease production, *ureABC*, but lacks any of the accessory genes (Fig. 1A). This results in very low intrinsic urease activity (29), providing a suitable background for genetic engineering. Previous work in *E. coli* had been based on heterologous expression of the urease gene cluster from *Sporosarcina pasteurii* as a means of driving MICP (17). We therefore initially focused on the same genes and could indeed impart enhanced urease activity to *B. subtilis* following inducible expression of the *S. pasteurii ureABCEFGD* operon (Additional File 1). Interestingly, while the intrinsic urease activity of the recipient strain, as expected, was below the detection limit of the colourimetric urease test broth used, we consistently observed that engineered urease activity was markedly higher when a chassis organism was used in which the native *ureABC* operon had been deleted (Additional File 1). This hinted at interference between the native and heterologous core proteins, and all further work was therefore carried out in a *ureABC* deletion background. While the results provided proof-of-concept evidence that heterologous urease gene expression could drive enhanced ureolytic activity in *B. subtilis*, these early experiments were beset with technical issues of strain stability and reproducibility of any quantitative data, indicative of toxicity of the introduced genes.

**Figure 1.**
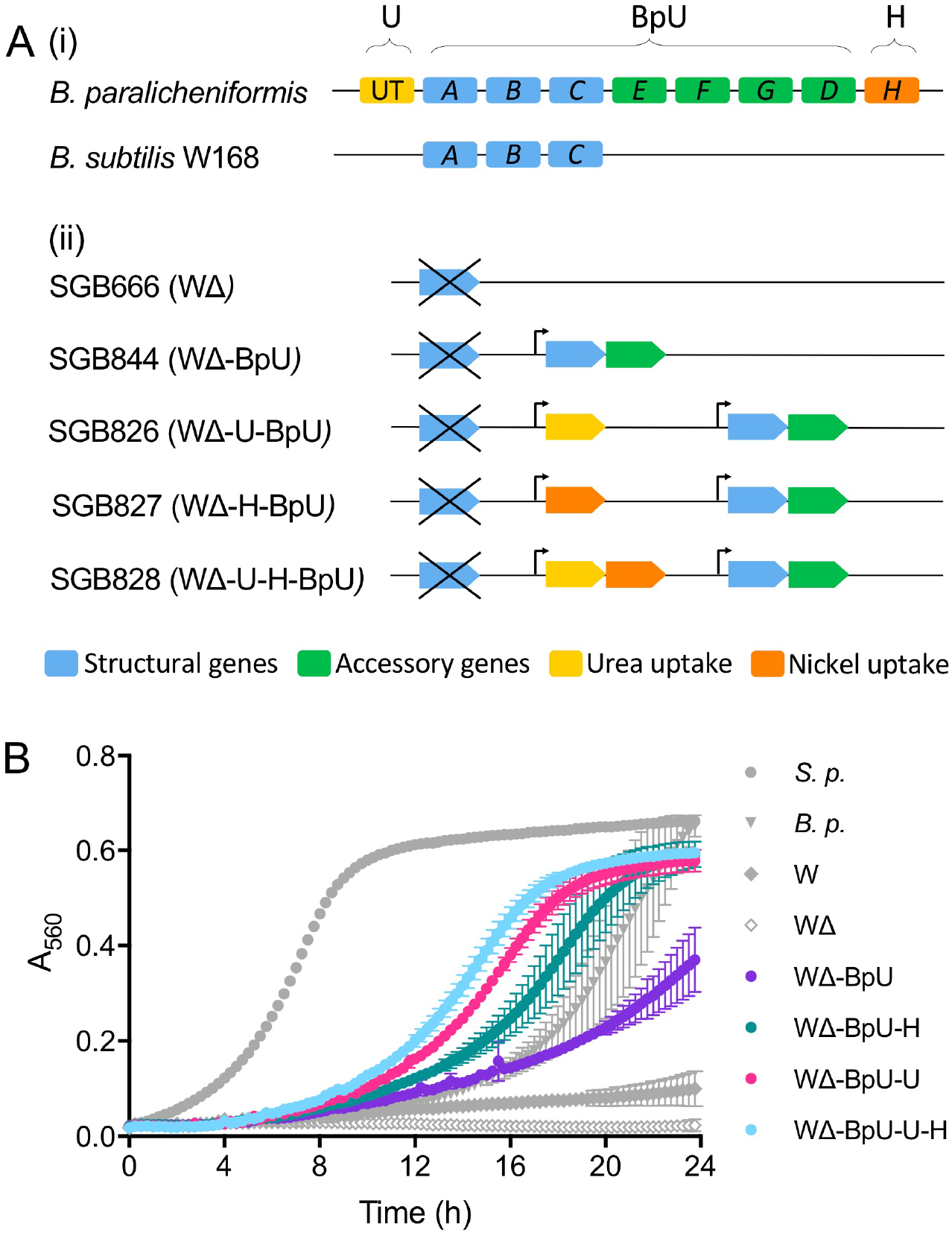
Heterologous expression of *B. paralicheniformis* urease genes in *B. subtilis*. **(A)** Schematic of urease gene loci. (i) shows the native urease genes of the donor and recipient species. Functional categories for each gene are indicated using the colour code given below the schematics. For simplicity, only the last letter of each gene name in the *ure* cluster is shown. The abbreviated terms used in strain denominations throughout this study is shown above. (ii) shows the construction of the heterologous expression strains. Deletion of the native *ureABC* operon is indicated by a crossed-out blue arrow. Introduced genes are shown in simplified schematics using the same colour code. Bent arrows indicate the P*_xylA_* promoter used to drive gene expression. **(B)** Urease activity of heterologous expression strains. Urease test broth containing 0.2 % xylose were inoculated to an initial OD600 of 0.05 from solid media growth and incubated at 30°C for 24 h in a TECAN Spark microplate reader. Absorbance at 560 nm (A_560_) was monitored to detect the yellow-to-pink colour change of the pH indicator phenol red as a result of urease activity. The composition of the test broth did not support cell growth and OD_600_ remained constant during the experiment, giving no interference with A_560_ measurements. The data are shown as mean ± standard deviation of 2-3 technical repeats and are representative of three independent repeats. Control strains shown in grey symbols are *S. pasteurii (S.p.), B. paralicheniformis (B.p.), B. subtilis* W168 (W), and its isogenic *ureABC* deletion (WΔ). The heterologous strains are shown in colour using the nomenclature detailed in panel A.

To improve the performance of the engineered strain, we therefore decided on a donor organism with a closer phylogenetic relationship to *B. subtilis. Bacillus paralicheniformis* ATCC9945a possesses a full urease gene cluster, harbouring the core genes *ureABC* as well as the accessory genes *ureEFGD*. Additionally, the gene cluster is flanked by a nickel permease, *ureH*, downstream of the urease genes, and a eukaryotic-type urease transporter, UT, upstream (Fig. 1A). This would allow us to test the effect of these additional genes during strain optimisation. Moreover, the *ureABCEFGDH* operon of *B. paralicheniformis* had been used successfully in the past to neutralise internal pH in a metabolically engineered *B. subtilis* background (30), giving confidence in its suitability for our purposes. To assess the relative contribution of the different genes to heterologous urease activity, a series of plasmids was constructed containing the main urease operon *ureABCEFGD* of *B. paralicheniformis* (‘BpU’); the urea transporter UT (‘U’); the nickel permease *ureH* (‘H’); or a synthetic operon of the two latter genes (‘U-H’). All of these were placed under control of the xylose-inducible promoter P*_xylA_*. These constructs were introduced into the *ureABC*-deletion background of *B. subtilis* W168 (WΔ) via stable, single-copy integration into the chromosome (Fig. 1A).

To compare urease activity between these strains, we adapted the standard qualitative colourimetric urease assay used initially to instead facilitate time-resolved detection of urease activity. In brief, the assay is based on a yellow-to-pink colour change of the pH indicator phenol red, responding to the pH increase resulting from the cleavage of one mole of urea to two moles of ammonia plus carbon dioxide, catalysed by urease. Using a microplate reader, we could track this colour change as an increase in absorbance at 560 nm wavelength (A_560_) over time, with higher urease activity seen as an earlier and steeper response (Fig. 1B). Incubation of *B. subtilis* W168 wild-type cells in this assay showed the expected very low intrinsic urease activity of this strain (Fig. 1B, grey diamonds), which was reduced to undetectable levels in the *ureABC* deletion strain (open diamonds). As a positive control, we included the strongly ureolytic *S. pasteurii* (grey circles), which showed a rapid colour change. The *B. paralicheniformis* donor strain of urease genes for our study likewise produced a clear colour change, but at an intermediate speed (grey triangles). These data showed the assay could accurately differentiate between levels of urease activity and produced no detectable background signal.

Heterologous expression in *B. subtilis* of the main urease gene cluster of *B. paralicheniformis* was sufficient to impart clearly detectable urease activity (Fig. 1B, purple symbols), although this did not reach the same activity as in the donor species. Additional expression of either the nickel (teal) or urea (pink) transporters drastically increased urease activity, exceeding the native activity of the donor strain. The urea transporter UT had the largest individual effect, while simultaneous expression of both transporters did not provide an additive benefit. Through heterologous expression of the *B. paralicheniformis* urease genes, particularly when co-expressed with the accessory transporters for nickel and/or urea, *B. subtilis* could therefore be engineered to display high ureolytic activity.

We next repeated these experiments on solid media, as growth on a surface would be more realistic for downstream application of MICP. For this, the strains were spotted onto LB agar supplemented with a calcium source (LBC) and the indicator phenol red. Media were prepared with and without addition of urea to allow the differentiation between genuine urease activity and low-level pH changes due to general metabolism of amino acids in the growth medium. pH changes in the agar were monitored photographically over 7 days, and urease activity scored as a stronger pH change compared to the same strain grown without urea. Under these conditions, we observed the same general trend as in the liquid test broth, where strong ureolytic activity required the inclusion of accessory transporters (Additional file 2A). Interestingly, on the richer LB medium, compared to the nutrient-poor colourimetric test broth, inclusion of the urea transporter, which previously showed the strongest individual effect, did not lead to a noticeable improvement. Instead, it was the nickel permease UreH that enabled the highest ureolytic activity. This showed that the two accessory transporters play different roles depending on the environment of the bacteria and whether access to nickel or urea is limiting to urease activity. Inclusion of both transporters consistently led to strong ureolytic activity in *B. subtilis* under both conditions. As the higher nutrient conditions in LBC were more representative of conditions used for the subsequent calcite precipitation assays, the strains co-producing urease with the nickel transporter (WΔ-H-BpU) and with both accessory transporters (WΔ-U-H-BpU) were taken forward.

### Heterologous production of urease in *Bacillus subtilis* W168 facilitates biomineralisation

To test if the strains with engineered urease activity had also gained the ability to precipitate calcite, we next spotted the two strains with strongest ureolytic activity onto precipitation media. For this, we first chose LBC medium as used for the urease activity assays, but without the pH indicator and initial adjustment of the pH to avoid potential interfering effects. After seven days of incubation, crystal formation was reproducibly visible on the colony surface for WΔ-H-BpU, but not for WΔ-U-H-BpU (Fig. 2A). While these results confirmed that introduction of urease activity could impart MICP activity to *B. subtilis*, it was surprising to see no significant crystal formation by strain WΔ-U-H-BpU, which had shown strong ureolytic activity in all previous assays.

**Figure 2.**
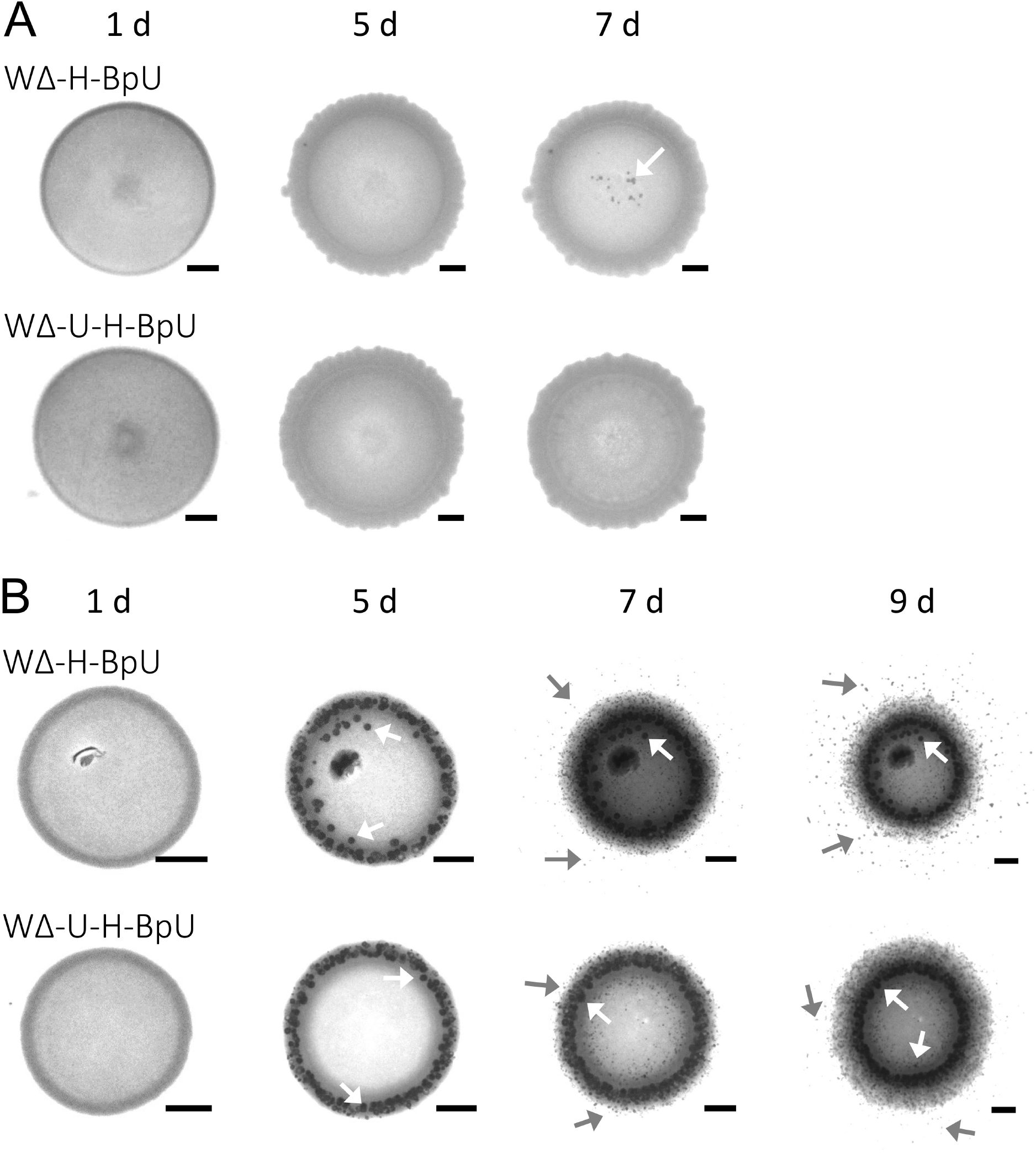
Mineral precipitation of urease-expressing strains of *B. subtilis*. **(A)** Precipitation on LBC media. **(B)** Precipitation on B4 media. Both media were supplemented with urea and xylose. Urease-producing strains WΔ-H-BpU and WΔ-U-H-BpU were spotted onto solid precipitation media and incubated at 30°C. Mineral precipitation was monitored for up to 9 days and images of the colonies taken with a stereomicroscope at the time points indicated above the panels. Mineral crystals formed on the colony surface are indicated with white arrows, crystals formed on the surrounding agar by grey arrows. Scale bars represent 1 mm in size. The results are a representative series from two to three biological repeats.

To address this further, we repeated the precipitation experiment using B4 agar, which is an established precipitation medium used to assess biomineralisation potential of environmental isolates (31). On this medium, precipitation onset was visible as early as three days for both WΔ-H-BpU and WΔ-U-H-BpU (Fig. 2B). This started with crystals localised to the colony surface and later spread to the surrounding media, visible clearly for both WΔ-H-BpU and WΔ-U-H-BpU after nine days, although crystal formation was again consistently more pronounced for WΔ-H-BpU. The earlier precipitation onset on B4 media was likely due to two reasons. Firstly, B4 medium has an initial pH of 8, while fresh LBC medium typically has a pH value around 7. Thus, in B4 medium, less ammonia production by urease is required to increase the pH to a level that is favourable for biomineralisation. Secondly, LBC contains considerably more complex components than B4, with the resulting buffering effect likely reducing the spread of the pH change through the medium and hence leading to crystal formation predominantly on the colony, rather than the surrounding agar. Given the favourable conditions for MICP on B4 media, both of the strongly ureolytic derivatives of *B. subtilis* had gained biomineralisation activity. Thus, heterologous expression of urease genes in *B. subtilis* can drive MICP, but the surrounding chemistry dictates the speed, spatial distribution and observable quantity of mineral production.

### The biomineralisation product from engineered *B. subtilis* W168 is calcium carbonate

To determine if the observed crystals were indeed the desired mineral calcium carbonate, a sample of crystals was removed from a colony of WΔ-H-BpU grown on LBC-phenol red agar and subjected to scanning electron microscopy (SEM) with energy dispersive X-ray (EDX) elemental analysis. SEM imaging revealed the harvested crystals to be aggregates of smaller crystals (Fig. 3A). In several of the analysed precipitates we observed smaller, round structures, which at higher magnifications were revealed as having a ‘dumbbell’ morphology and appeared embedded in a web-like matrix, which may be indicative of organic material in the crystal aggregates (Fig. 3C&D). This dumbbell morphology was frequently observed in biological repeats of the experiment, but not always to the same extent, suggesting that the formation of these dumbbell structures was sensitive to conditions and timing of sampling. Crystals with this shape have been observed previously in association with microbial activity (32, 33). Although definitive experimental evidence for this is missing, it is possible that encased bacteria lie at the centre around which layers of mineral are deposited over time until they turn from a dumbbell morphology into larger spherical morphologies (Fig. 3C), as has been reported for bacterial dolomite precipitation (32).

**Figure 3.**
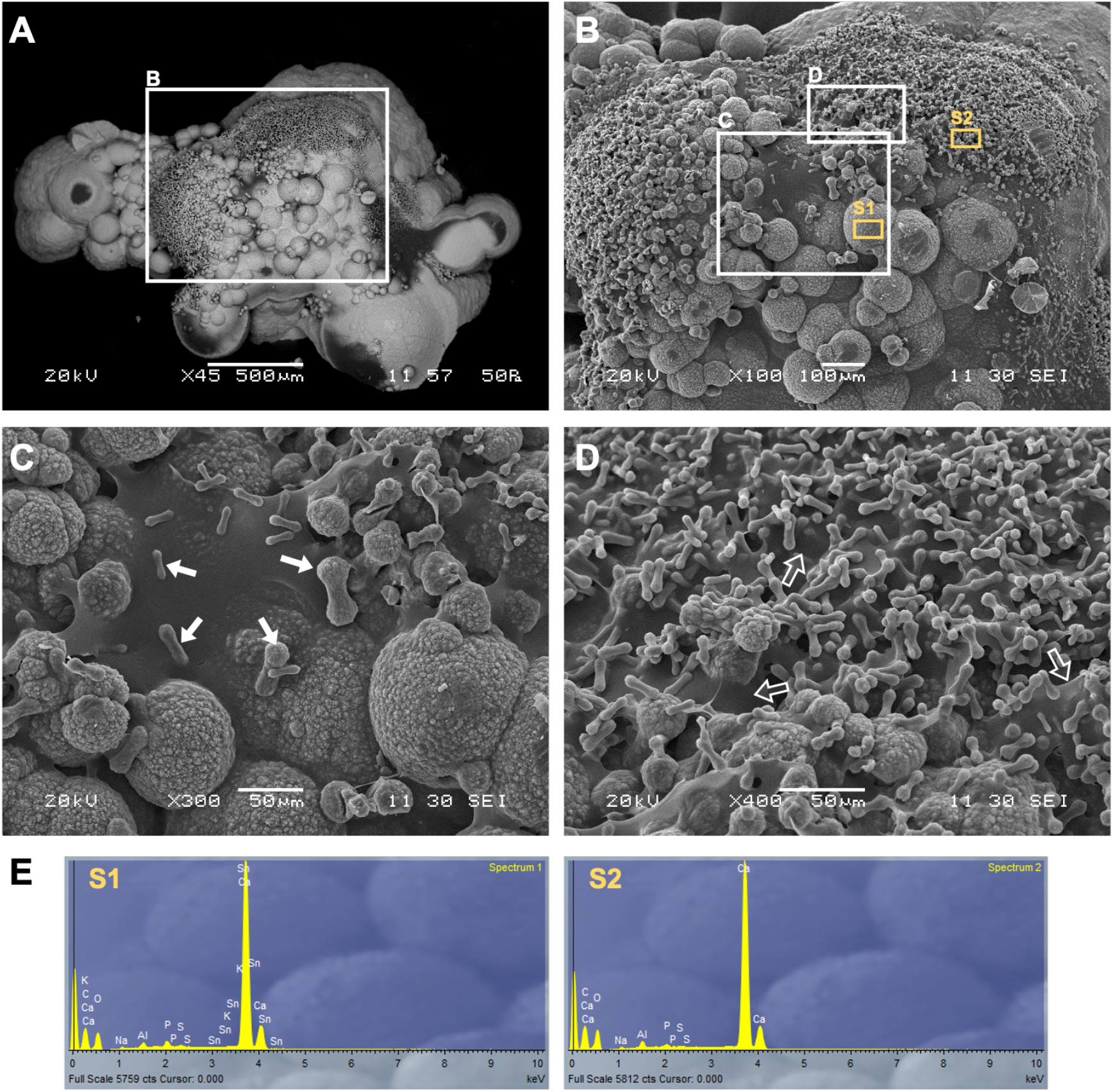
Scanning electron microscopy of calcite crystals produced by strain WΔ-H-BpU. Mineral crystals formed after incubation of strain WΔ-H-BpU on LBC-phenol red agar for two weeks at 30°C and one week at room temperature were removed from the colony surface and imaged by scanning electron microscopy with gold splutter coating. **(A-D)** Sequential zoom in on a crystal particle. The white boxes in panels A and B indicate areas of view shown in the subsequent panels. White arrows indicate the different sized stages of crystals in the commonly observed ‘dumbbell’ morphology. Open arrows indicate the web-like material observed between the dumbbell shaped crystals. **(E)** EDX elemental analysis was performed at two image sites (S1 and S2) indicated in panel C to verify the precipitate composition was predominated by calcium (Ca), carbon (C) and oxygen (O).

Analysis of the elemental composition of the crystals via EDX spectroscopy revealed that the precipitates were indeed calcium carbonate, as evidenced by the predominant peaks for Ca, C and O (Fig. 3E). While insufficient material was produced by our experiments to allow the identification of the specific polymorph, the crystals are most likely formed of calcite, as this is the most stable and abundant form of the mineral (34).

### Biofilm-promoting conditions enhance biomineralisation in *B. subtilis*

Our next aim was to study the impact of biofilm formation on biomineralisation, because the extracellular matrix can aid mineral precipitation by creating a suitable microenvironment, e.g. through trapping of ions or provision of nucleation sites (5). As *B. subtilis* strain W168 is an intrinsically poor biofilm former, we instead turned to the strong biofilm producer strain *B. subtilis* NCIB3610. To establish a basal level of biomineralisation activity, we first reconstructed heterologous urease production as described above in this new strain background. This gave essentially the same results as in strain W168, with strains NΔ-H-BpU or NΔ-U-H-BpU (NΔ denoting the native *ureABC* deletion in strain NCIB3610) giving the highest urease activity on LBC medium with urea and phenol red (Additional file 3A). Precipitation results were also comparable, although the NCIB-derived strains had even lower precipitation ability on LBC medium than the W168-derived strains, with neither NΔ-H-BpU nor NΔ-U-H-BpU producing visible crystals after 9 days. On B4 medium, both of these strains showed strong biomineralisation activity, again with NΔ-H-BpU giving the strongest precipitation on the colony surface and the surrounding agar (Additional file 3B&C). This showed that the heterologously expressed urease genes bring out similar phenotypes across *B. subtilis* strains.

To induce biofilm formation, we initially used MSgg medium in line with standard protocols for *B. subtilis* biofilm growth (35). However, after addition of urea and calcium to facilitate precipitation, growth of our engineered strains on this medium was poor. We therefore turned to the complex biofilm promoting medium LBGM (LB medium supplemented with glycerol and manganese) (36), which supported robust growth after addition of urea and calcium (LBGMC). It was also most similar to the LBC medium used in the earlier experiments and would provide a good basis for comparison of the data. When strains NΔ-H-BpU nor NΔ-U-H-BpU were spotted onto this medium, they showed a clear biofilm morphology with growth over a much larger surface area. Inspection for biomineralisation revealed the presence of numerous crystals after seven days of incubation (Additional file 3D). Compared to the lack of precipitation seen on plain LBC medium, this clearly showed that biofilm formation had an enhancing effect on biomineralisation. However, the major changes in the colony morphology of the NCIB3610-derived strains made it difficult to accurately quantify the differences. Because the increased surface area provided more potential sites for precipitation, it was unclear whether the enhanced precipitation was the result of the larger colony area or of biofilm formation *per se*.

To more precisely establish the links between biofilm formation and mineral precipitation, further testing was required in a strain that did not undergo major morphological changes on LBGMC medium. We therefore returned to the W168-derived strains, which showed colonies of similar size and structure between LBC and LBGMC media. Monitoring crystal formation of strain WΔ-H-BpU following spot-inoculation onto both media revealed a striking increase in the number of crystals formed under biofilm-promoting conditions. Onset of appearance of visible crystals was also much earlier, with multiple crystals visible after three days, compared to seven days on LBC agar (Fig. 4). Thus, while W168 is not traditionally the strain of choice to study biofilm formation, growth under biofilm-promoting conditions did clearly enhance biomineralisation. Moreover, the uniform colony morphology of the W168-derived strains enabled accurate enumeration of crystals and provided a suitable experimental system to take forward to elucidate the contribution of individual elements of biofilm formation on calcium carbonate precipitation. It should be noted that the WΔ parent strain without urease expression never formed any visible crystals when grown on LBGMC medium for up to ten days. Biofilm-promoting conditions alone were therefore not sufficient to drive biomineralisation in the absence of urease activity.

**Figure 4.**
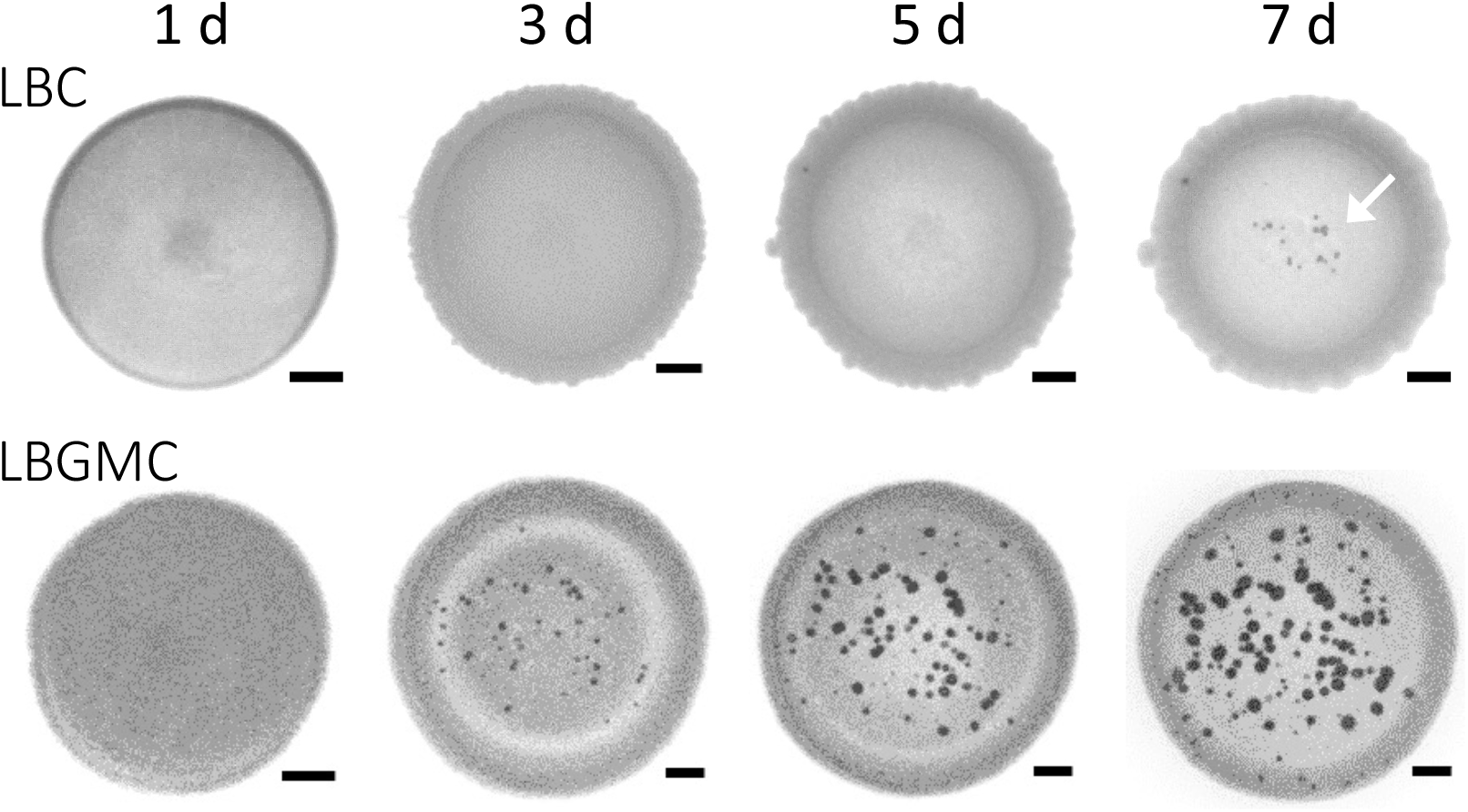
Effects of biofilm formation on biomineralisation by *B. subtilis*. Cells of strain WΔ-H-BpU were inoculated into the centre of LBC agar plates or on biofilm-promoting LBGMC plates. Both media were supplemented with urea and xylose. Mineral precipitation was monitored over seven days of incubation at 30°C and images of the colony taken under a stereomicroscope at the time points indicated above. Scale bars represent 1 mm in size. The results shown are a representative series from three to four biological repeats. The images at days 1, 5 and 7 on LBC medium are the same as shown in figure 2.

### The polysaccharide but not protein component of biofilms drives biomineralisation

Following the observation that biofilm-promoting conditions greatly enhanced mineral formation, we next aimed to identify which component of the biofilm was responsible for this effect. *B. subtilis* biofilm compositions involve extracellular polymeric substances which are made up of polysaccharides, charged polymers, amphiphilic molecules and proteins, where the major components are polysaccharides and proteins (24). To differentiate between the role of the polysaccharide versus proteinaceous component of the biofilm matrix, we targeted two biosynthetic genes, *epsH* and *tasA*. The former encodes a glycosyltransferase for exopolysaccharide synthesis, while the latter encodes the extracellular fiber forming protein TasA (24, 37). Both genes have been shown to be upregulated on LGBM media ensuring expression under our experimental conditions (36). To test their role in calcite precipitation, the *epsH* and *tasA* deletions were each introduced into the strong precipitator WΔ-H-BpU. This led to the construction of strain WΔ-Δe-H-BpU and strain WΔ-Δt-H-BpU, respectively.

When examining the biomineralisation of the *tasA* deletion strain compared to its WΔ-H-BpU parent strain no clear difference in precipitation was exhibited following growth on LBGMC medium over five days at 30°C (Fig. 5, open squares). This was supported by statistical analysis using a standard two-way ANOVA test, which showed a significant effect of time (p<0.0001), but not of strain background (p=0.8610) on the number of observed crystals. However, comparison of the *epsH* deletion strain to the WΔ-H-BpU control, revealed a clear decrease in the amount of precipitation on the colony (Fig. 5, grey circles). A two-way ANOVA confirmed a significant effect of time (p=0.0001), as well as strain background (P=0.0005). This showed that the precipitation-promoting effect of biofilm formation was specifically attributable to the polysaccharide component of the biofilm matrix, while the TasA protein did not appear to contribute.

**Figure 5.**
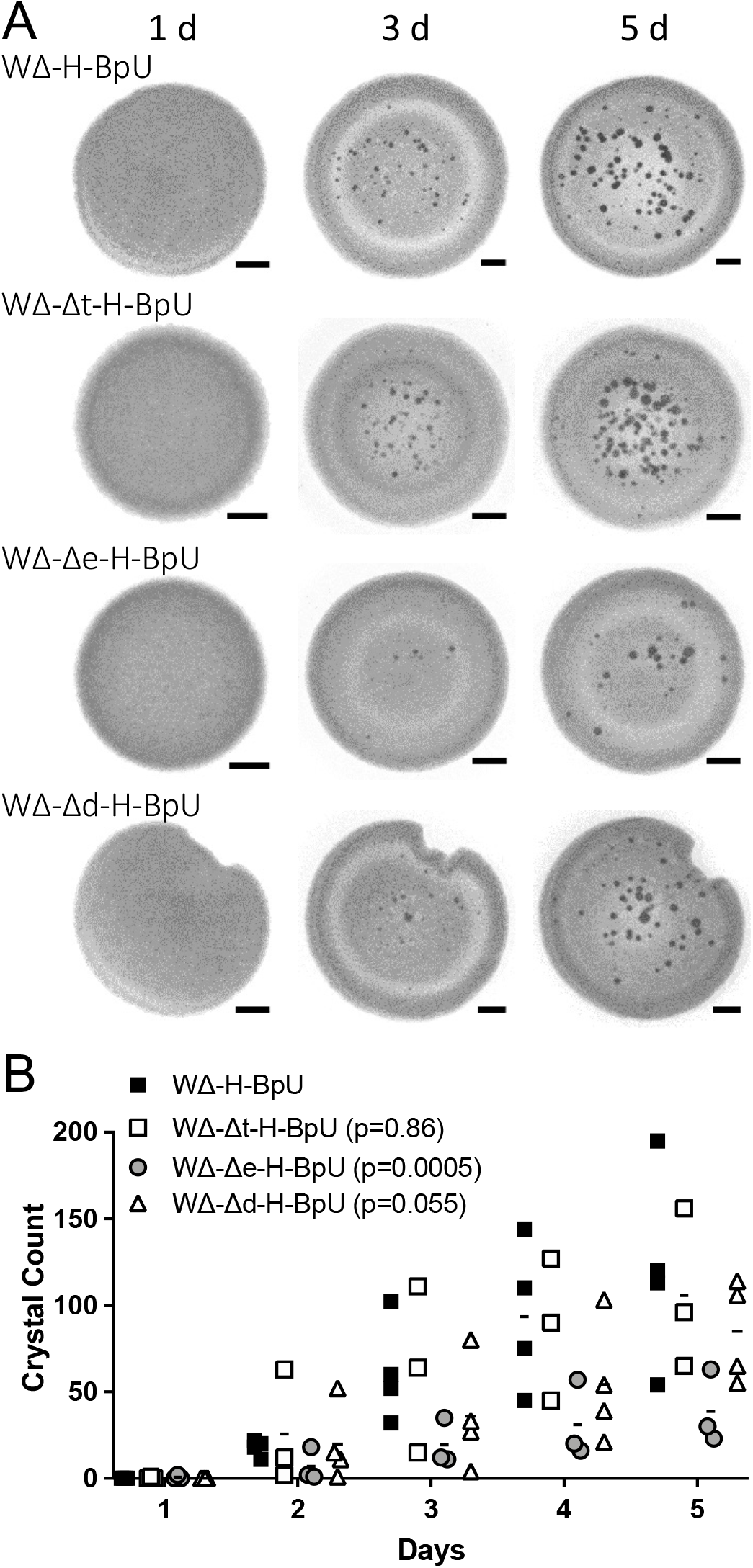
Contribution of biofilm components and surface charge to biomineralisation. **(A)** Crystal formation on LBGMC agar. Cells of the indicated strains were inoculated into the centre of biofilm-promoting agar plates containing urea and xylose and monitored over five days of incubation at 30°C with images of the colony taken under a stereomicroscope at the time points indicated above. Scale bars represent 1 mm in size. The results shown are a representative series from three to four biological repeats. The images shown for WΔ-H-BpU are the same as shown in figure 4 and serve as the baseline control. **(B)** Quantification of crystal formation (crystal count per colony) over time. Individual data points for all biological repeats are shown, with the average value depicted as a line. The inset legend gives the p-values for differences by strain from a two-way ANOVA for each dataset compared to the WΔ-H-BpU parent strain. Δt, *tasA* deletion; Δe, *epsH* deletion; Δd, *dltABCDE* deletion; remaining strain nomenclature as in previous figures.

### Manipulation of the cell surface charge via *dltABCDE* deletion does not impact biomineralisation in *B. subtilis*

Cell surface charge is thought to play a key role in mineral precipitation where the net negative bacterial surface attracts metal cations, and the resulting locally increased concentration favours precipitation (reviewed in (5)). In Gram-positive bacteria, the surface charge is modulated by the action of the Dlt system, encoded by *dltABCDE*, which reduces the net negative charge via D-alanylation of teichoic acids. In the absence of Dlt the cell surface is expected to bear a greater negative charge due to the decreased masking of phosphate backbone negative charges by positively charged amine groups. To test if this could be exploited to increase the attraction of calcium ions and thus calcium carbonate precipitation in *B. subtilis*, we deleted the *dltABCDE* operon of strain W168, resulting in strain W-Δd. Surprisingly, zeta potential measurements of the deletion strain showed a decrease in negative potential from -61 mV to -57 mV compared to wildtype instead of the expected change towards a more negative charge (Additional file 4A). To exclude that this was due to the assay used, we next tested both strains for binding of cytochrome C, which is commonly used to characterise surface charge whereby the adsorption of the positively charged cytochrome C protein to bacterial cells is used as a proxy for negative surface charge. Here, we observed a marginal, but not statistically significant increase in binding from 67% in the wild-type to 72% in the *dlt* deletion strain (Additional file 4B). This small change was in stark contrast to the effect of *dlt* deletion in other Gram-positive bacteria, where a change from approximately 10% to 70% of cytochrome C binding has been reported upon *dlt* deletion in *Staphylococcus aureus*, and a change from 30% to over 80% binding in *Lactobacillus casei* (38, 39). To exclude any issues with our *dlt* deletion strain, we next tested its sensitivity to the cationic antimicrobial peptide nisin. The Dlt-mediated reduction in the net negative surface charge has been reported to contribute to resistance, and a previously described *dlt* deletion mutant of *B. subtilis* showed increased nisin sensitivity (23). When cultures of our wild-type and *dlt* deletion strain were challenged with increasing concentrations of nisin, the wild-type strain showed growth inhibition upon addition of 32 μg/ml nisin, while the *dlt* deletion strain showed a similar degree of growth inhibition already at 16 μg/ml (Additional file 4C). This confirmed that our deletion strain did indeed display expected differences to its parent strain, but the overall effect on net surface charge appeared to be unexpectedly minor.

To test if *dlt* deletion could nevertheless improve calcium carbonate precipitation, we introduced the deletion into the strong precipitator background described above, resulting in strain WΔ-Δd-H-BpU. When this strain was subjected to the same precipitation assay used before, it showed similarly strong biomineralisation as the parent strain WΔ-H-BpU, and we did not detect a significant change in either the time to onset of precipitation or the number of crystals formed (Fig. 5). The lack of an obvious impact on precipitation suggests that the Dlt-dependent modulation of surface charge in *B. subtilis* was too minor to have a measurable effect and/or the surface charge of this species was already sufficiently negative to enable biomineralisation.

### Biofilm-promoting conditions create layered structures within precipitated minerals

To investigate the effects of biofilm formation on the resulting mineral precipitate, we subjected crystals lifted from the bacterial colonies grown on LBGMC agar to SEM analysis. Compared to the fairly simple aggregated crystals observed under standard conditions shown above, the crystals from the biofilm-promoting conditions showed a striking layered morphology (Fig. 6A&B). Areas more central to the harvested particles showed an open, porous structure, whereas areas at the particle surface showed a smooth layer covering the porous material. This suggests that under these conditions, the mineral precipitation mostly occurs on or within the extracellular matrix of the biofilm, consistent with the contribution of the polysaccharide component to precipitation, reported above, leading to the smooth appearance. The porous central sections may be explained by holes originating from the spaces originally taken up by bacterial cells. This is supported by some fields of view showing very clear ‘footprints’ of bacterial cells, where the mineral was precipitated around the cells and the imprint of the cell remained visible as negative space (Fig. 6C). Together, our findings indicate a sequential formation of layers over time during biomineralisation under biofilm-promoting conditions, with the bacteria themselves at the centre resulting in a porous precipitate, surrounded by a smoother layer where precipitation occurred predominantly in the extracellular matrix. Consistent with our earlier data, we could not observe consistent differences between the strong precipitator strain WΔ-H-BpU and its *dlt*-deletion derivative (Fig. 5), further supporting our conclusion that in *B. subtilis* this pathway of surface modification, under the conditions tested here, does not significantly contribute to enhancing biomineralisation.

**Figure 6.**
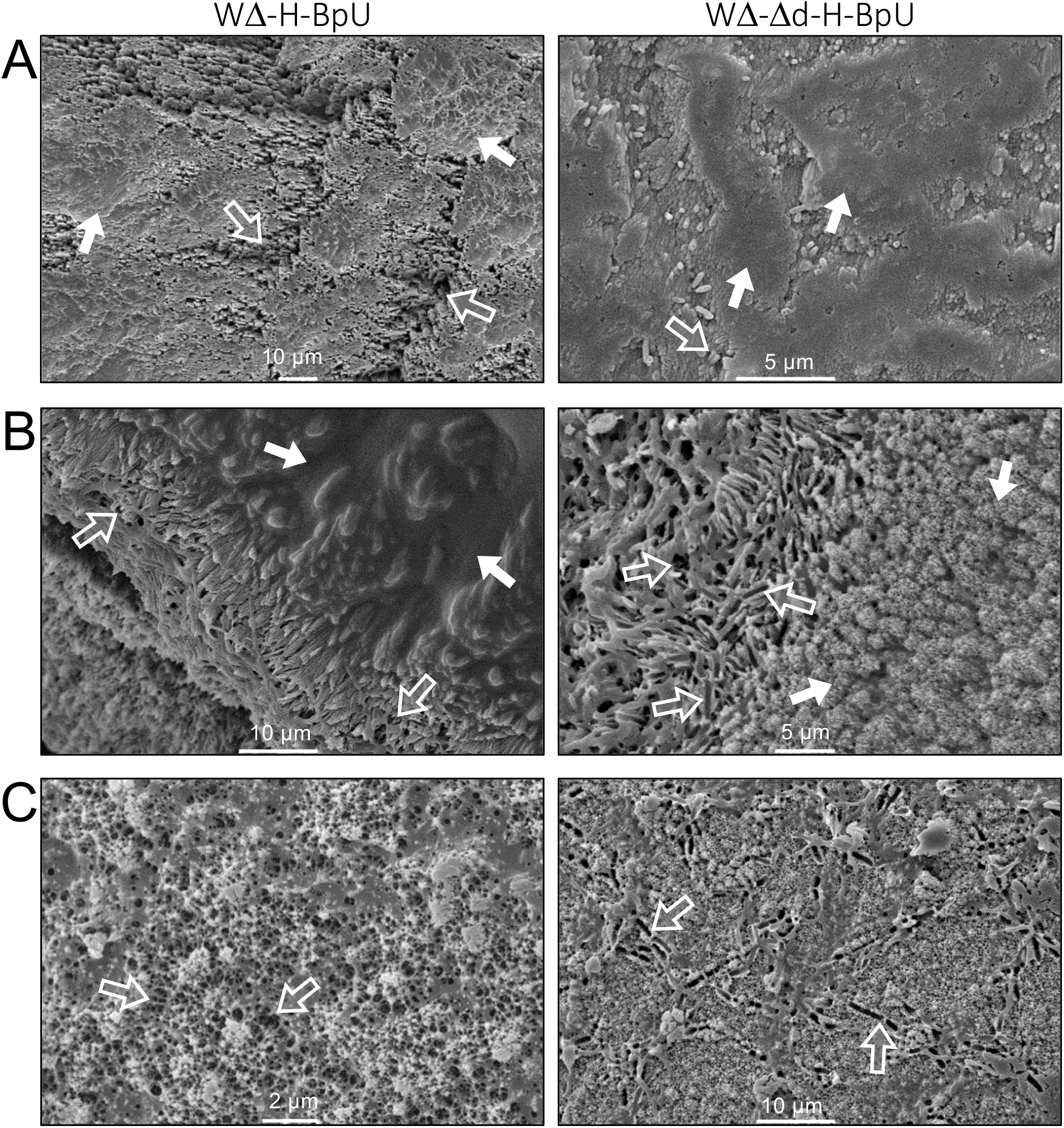
Scanning electron microscopy of mineral precipitates formed under biofilm-promoting conditions. Crystals precipitated on colonies of WΔ-H-BpU (left panels) and WΔ-Δd-H-BpU (right panels) were grown on LBGMC agar containing urea and xylose at 30°C for 1-2 weeks. Crystals removed from the colony surface were imaged on an SEM with gold splutter coating. **(A)** Areas of view showing the particle surface with porous sections visibly covered by a smooth layer. **(B)** Areas of view showing boundaries between smooth and porous sections of the precipitate. **(C)** Views of porous sections of the precipitate. Closed arrows indicate smooth areas of precipitate, presumably within the extracellular biofilm matrix; open arrows indicate porous areas of precipitate and voids or bacterial ‘footprints’ in the precipitate. Sizes of scale bars are given in each image.

## Discussion

The aim of this investigation was to use a bottom-up approach to engineer the Gram-positive bacterium *B. subtilis* W168 for MICP to understand which bacterial and genetic features or processes drive biomineralisation and how their interactions may lead to optimal precipitation. This was approached by exploring the role of ureolysis, biofilm formation, and cell surface charge on biomineralisation, which showed that maximum MICP resulted from production of urease in conjunction with the nickel permease UreH and growth under biofilm promoting conditions, while the activity of the Dlt system did not have significant effects. A diagram summarising the multiple contributing factors in the engineered precipitator strain is presented in Figure 7.

**Figure 7.**
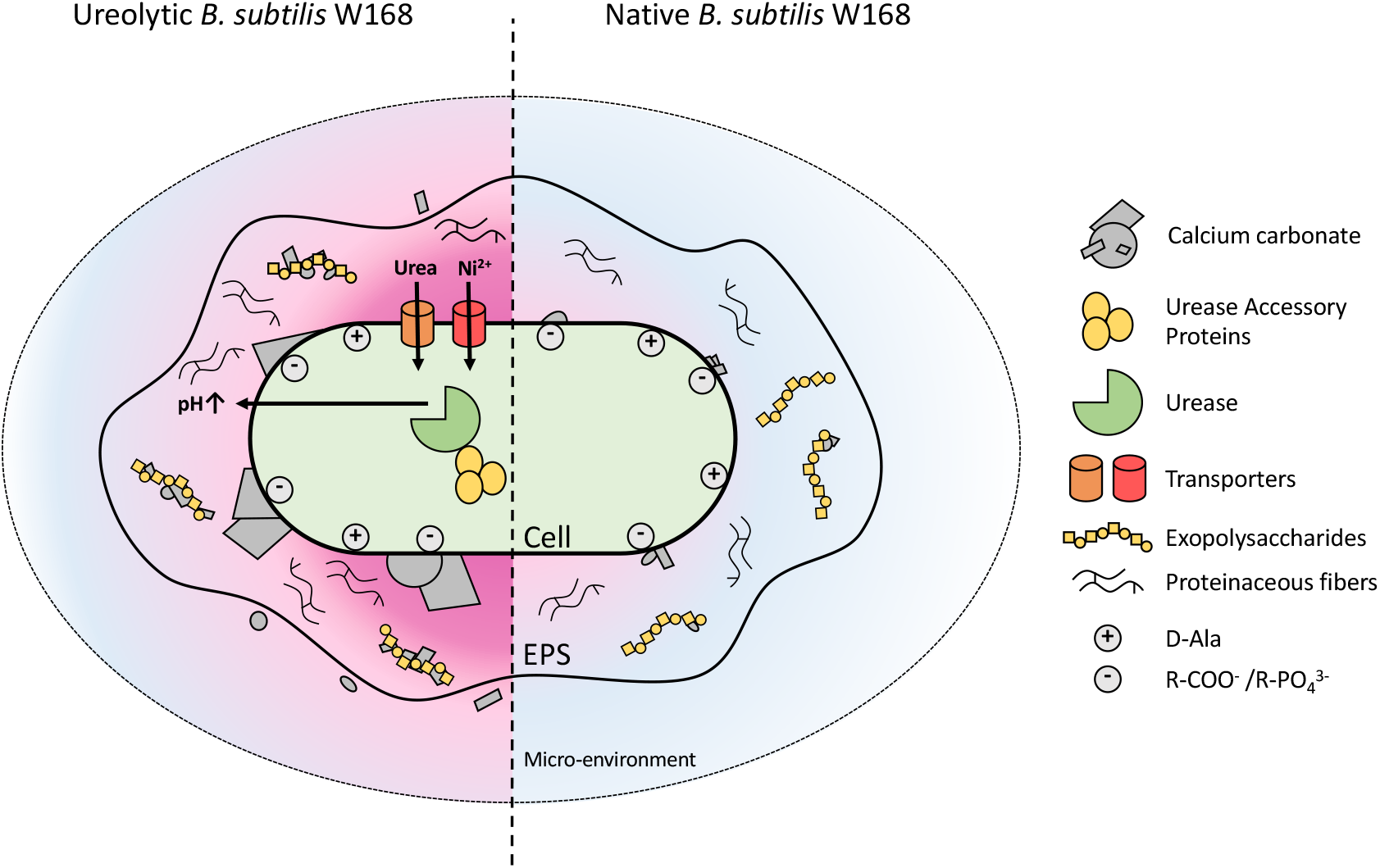
Schematic diagram of engineered calcium carbonate precipitation in *Bacillus subtilis* W168. The cellular processes and features are shown that drive biomineralisation in the engineered ureolytic precipitator strain (left) and native *B. subtilis* W168 (right). In the precipitator strain, heterologously produced urease catalyses the breakdown of urea leading to an increased pH of the microenvironment (pink gradient) which favours calcium carbonate precipitation. The presence of accessory proteins UreE, UreF, UreG, and UreD as well as nickel and urea transporters contribute to increased ureolysis and precipitation. At sufficiently high pH, precipitates may even form in the micro-environment surrounding the cell. The presence of exopolysaccharides within the biofilm matrix amplifies precipitation promoted by the increased pH and may act as additional nucleation templates. Proteinaceous fibres were not found to promote precipitation. Negatively charged sites on the cell surface likely act as nucleation sites in both native and engineered *B. subtilis*, but are by themselves insufficient to drive detectable biomineralisation. Native *B. subtilis* W168 increases the localised pH only slightly from metabolic activities (such as amino acid degradation), which is not sufficient to reach conditions required for precipitation. Low levels of undetected precipitates are still likely to form on the cell surface and extracellular matrix.

Previous studies of engineered MICP via ureolytic activity predominantly focussed on the urease gene cluster of *Sporosarcina pasteurii*, the prototypical species for application in MICP-based biotechnology. These studies showed that heterologous expression of the genes could impart ureolytic and MICP activity to *E. coli* (17–19). While we here showed that the same genes could also be used to introduce urease activity to *B. subtilis*, we obtained more robust results with *B. paralicheniformis* as the donor species for the gene cluster, indicating differences in genetic compatibility between donor and recipient. Early studies of the genetic determinants of urease activity had also revealed that the requirement for specific urease genes was context dependent: a sub-set of urease genes of *Helicobacter pylori* could impart ureolytic activity when heterologously expressed in *Campylobacter jejuni*, while activity in *E. coli* required co-expression with further accessory genes (40, 41). These findings highlight the need to study heterologous urease production in application-relevant organisms to account for such variations between donor and host compatibility.

Moreover, our choice of donor species allowed us to systematically test the contribution of accessory transporters for nickel and urea associated with the urease operon of *B. paralicheniformis*. Our results showed that co-expression of the accessory transporters was generally beneficial to the urease activity of the resulting *B. subtilis* strain, but revealed striking differences depending on the conditions under which the activity was assessed: under low-nutrient conditions in the qualitative test broth, urea uptake had the most significant individual effect; while under nutrient-rich conditions in LB-based media, nickel uptake was the most relevant. Importantly, the speed of mineral precipitation by the engineered strains correlated well with their respective strength of ureolytic activity, with strains WΔ-H-BpU and WΔ-U-H-BpU, carrying the core urease genes plus the accessory nickel transporter gene *ureH* alone or together with the urea transporter UT, showing the most rapid and most extensive crystal formation. This shows that ureolysis can be an effective driver of MICP in *B. subtilis*, and that rapid assessment of ureolytic activity can be used as a good predictor of biomineralisation potential, facilitating time-effective strain optimisation.

In addition to the varied strength of ureolytic and MICP activity depending on the gene set expressed, we also noticed a strong impact of media composition on biomineralisation. Precipitation on complex medium (LBC) was comparatively slow and only occurred on the colony surface, while on minimal medium (B4) it was much faster with crystals forming on the colonies as well as on the surrounding agar. This is likely due to the differences in buffering capacity, which is considerably lower in B4 medium and would thus allow the optimal pH and saturation conditions for calcium carbonate precipitation to be reached faster and with wider spread. The effect of media composition on MICP has not been subject to much systematic study in the laboratory, although data from materials science investigations have shown that media composition, including presence of agarose, will affect solute movement and thus the supersaturation state required for mineral precipitation (42, 43). Such effects should be kept in mind when designing MICP strains for application. While urease activity can be a clear driver for biomineralisation, the surrounding chemistry is a key factor in determining the exact outcome.

To fully exploit the information gained here under laboratory conditions, it is therefore important to factor in the conditions likely encountered during application. In terms of the optimal gene set for ureolytic activity, it is plausible that nickel would be more available in applications, e.g. in water, soil or cementitious materials, than under controlled laboratory conditions using distilled water in the growth media. Hence the relative importance of the nickel transporter UreH identified as most beneficial in our study under LB-based conditions may prove to be reduced in some applied settings. We showed that under nutrient-poor conditions, accessory urea uptake enhanced biomineralisation, and this may have direct relevance to many applications where the nutrient provision will likely be limited. Indeed, it has been shown in a self-healing concrete system that the quantity of yeast extract provided correlated with the speed of biomineralisation, showing that under standard conditions self-healing was limited by access of the bacteria to nutrients (44). To give most robust performance, a strain harbouring both urea and nickel transporters may be ideal to cover both limitations. However, our results showed that strain WΔ-U-H-BpU produced less calcite than WΔ-H-BpU, possibly due to increased demand on the cell by production of an additional foreign protein. Further studies to improve the genetic compatibility and minimise energetic burden on the chassis organism may solve such problems and could lead to the development of a universally applicable, optimised MICP strain.

While in the present study we mainly focussed on the UreH-producing strains, it will be worth exploring the relative importance of the UT transporter in application settings. Past studies into the impact of urea uptake on ureolytic activity of bacteria have been centred on the high-affinity UrtABCDE system, encoded by the five-gene *urt* operon (13–15). The donor species we chose here, *B. paralicheniformis*, instead harbours a eukaryotic-type low affinity urea uptake system, UT. The fact that this transporter is encoded by a single gene might make it an attractive tool for enhancing ureolytic activity through genetic engineering under conditions where urea uptake is limiting.

At first glance it appears that strain optimisation should simply aim for stronger ureolytic activity, driving a faster pH change and thus leading to earlier biomineralisation. However, increasing evidence is emerging that the speed of MICP has a direct influence on the properties of the resulting mineral. For example, modulating urease activity in an *E. coli* strain harbouring the *S. pasteurii* urease genes via changes in gene copy number and the ribosome binding sites showed that ureolytic strength had a negative relationship with crystal size (17, 18). Comparison of different environmental strains showed that those with lower ureolytic activity induced localised precipitation on or near the bacterial cells, while strains with higher activity caused wide-spread precipitation in the media surrounding the bacteria (45). This is reminiscent of the media effects we observed here, showing that similar effects can be caused by either the media composition or the strength of the microbial activity. Simply increasing ureolytic activity may thus not be favourable for applications requiring larger crystals or controlled spatial localisation of the precipitate, where more delicate fine-tuning of MICP may be an advantage.

Biofilm formation is well-recognised to influence MICP, although its precise contributions and the underpinning molecular mechanisms have remained elusive. This is likely due to the complexity of both the biofilm matrix and the MICP process. For example, the extracellular polymeric substance of a biofilm can have either a mineralizing or a dissolving effect, depending on its composition. Functional groups that attract carbonate and calcium ions can act as nucleation sites for crystal formation, but excessive chelation of calcium ions, e.g. via acidic monomers, can inhibit biomineralisation by sequestering the cations (9). Understanding these chemical processes in application-relevant microorganisms may provide a means of controlling and manipulating the MICP process. Moreover, biofilm formation potentially can be used as a structural element to target mineral deposition spatially through controlling biofilm growth (10), adding further relevance to a deeper understanding of the interplay between biofilm formation and biomineralisation.

Genetic engineering approaches at gaining such an understanding are challenging due to difficulties in separating causes and effects in the biofilm-MICP interactions. For example, earlier studies in *B. subtilis* NCIB3610 have reported that *epsH* and *tasA* deletions negatively affected the localization of calcite crystals within the colony architecture (28). However, the deletion strains by themselves show a loss of structure in their biofilm architecture, preventing clear conclusions on the actual cause for the impaired biomineralisation. We here met similar issues when studying the NCIB3610-derived MICP strains, which showed strongly elevated biomineralisation under biofilm-promoting conditions, but this may at least in part be explained by the increased surface area of the colonies providing more nucleation sites. To circumvent this inherent complexity of multiple interconnected contributors to MICP, we therefore based our investigation on *B. subtilis* W168, which does not overtly change its colony morphology under biofilm promoting conditions, allowing us to separate colony structure from composition of the extracellular biofilm matrix. These experiments allowed us to conclude that the fibre-forming protein TasA does not, in fact, appear to have a beneficial effect on biomineralisation, but the polysaccharides produced by the *eps*-operon encoded proteins promoted crystal formation.

Interestingly, this affect appears to be dependent on growth conditions and potentially genetic background. An earlier study on *Bacillus* sp. JH7 showed that growth on glycerol could stimulate exopolysaccharide production, but this prevented, rather than stimulated MICP (46). This again highlights the complexity of the MICP process, where excess chelation of calcium by monosaccharides and/or acidification of the micro-environment from metabolism of available carbohydrates can have unintended negative impacts on biomineralisation (9, 46). The interplay between surface or extracellular chemistry and pH is also clear from the results reported here, where we could only observe MICP, even under biofilm-promoting conditions, when the bacteria had the additional ability to increase the local pH via heterologous urease production. Thus, not only the properties of the bacteria themselves, but also the environment and nutrients they are provided with have to be carefully considered when engineering bacteria for MICP and designing technological applications. For example, when considering application in self-healing concrete, the pH of the surrounding environment is already very alkaline, so a strong metabolic drive for further pH increase may be less important than production of suitable exopolysaccharide to promote MICP. Our findings that the *eps* operon of *B. subtilis* W168 encodes the production of such exopolysaccharide with beneficial effects on biomineralisation may pave the way for targeted engineering of biofilms with enhanced MICP properties through controlling not only where a biofilm grows, but also how its extracellular polymeric substance is composed.

The final property we investigated for its effect on MICP was the cellular surface charge, which we attempted to modify by deletion of the *dlt* operon to prevent D-alanylation of teichoic acids. Surprisingly, characterisation of the resulting strain revealed only very minor apparent changes to the cell’s surface, detectable only in an assay testing for sensitivity to killing by the cationic antimicrobial nisin. Likewise, no significant change of MICP activity was observed upon *dlt* deletion. Given the minor effect on surface properties by the deletion, it is likely that the change in charge was too subtle to have a measureable effect. However, it has also been proposed that in ureolytic bacteria, exemplified by *S. pasteurii*, the notion of negative surface charge being important for cation attraction for MICP could be an oversimplification, with metabolic activity and urea cleavage being the main driver (47). Our data could be seen as consistent with such a model.

Interestingly, our results revealed that there are large differences in the relative contribution of the Dlt system to surface charge modification between Gram-positive bacteria. As stated earlier, the *B. subtilis* wild-type strain showed a much higher degree of cytochrome C binding compared to wild-type *S. aureus* and *L. casei*, in fact reaching similar values to the *dlt* deletion strains of those other species (38, 39). This may suggest that in *B. subtilis* the Dlt system plays only a comparatively minor role in charge modification and the bacterium intrinsically carries a greater net negative surface charge than some other Gram-positive species. It has been suggested that a strong negative zeta potential may be the explanation for the ability of *Bacillus sphaericus* to colonise limestone, which carries a positive zeta potential (48). If an intrinsically low zeta potential is a common property among *Bacillus* species, this may offer an explanation for their frequent use in MICP-based technologies. Further engineering of the surface chemistry may, in this case, not be required in developing an optimal precipitator strain.

## Conclusions

Taken together, our results showed that heterologous urease expression was sufficient to bring about a precipitation phenotype in a bacterial strain of low intrinsic MICP ability. This precipitation was dependent on the strength of ureolytic activity that could be enhanced by the supply of accessory nickel (*ureH)* and urease (UT) transporter genes. Thus, bacterial metabolic activity can be a driver of bacteria-induced precipitation. The more strongly this metabolic activity modulates supersaturation conditions, the more likely precipitation will occur. The findings present a systematic analysis of variations in urease gene composition on whole cell urease activity and calcite precipitation of a Gram-positive model and industrially relevant organism. Furthermore, our results showed that biofilm production synergises with ureolytic activity to increase the calcite precipitation, but was not on its own sufficient to drive biomineralisation. Of the biofilm components, exopolysaccharides were found to be the largest contributors to precipitation. Lastly, the results here also showed that modulation of surface charge was insufficient to promote precipitation in *B. subtilis* W168. From this comprehensive investigation of molecular features that drive MICP, the genetic targets *ureABCEFGD*, *ureH*, UT and *epsH* showed the best potential to control precipitation in *B. subtilis*. Heterologous expression and changes in regulation of these genes in synergy should provide optimal targets for future engineering of chassis organisms that are better suited to conditions in applied settings into strong precipitators for MICP-based biotechnologies.

Our findings also highlighted mutual influences between genetic complement of the bacteria on the one hand and composition of the media or environment on the other in determining speed, spatial distribution and crystal properties of MICP. This means that true synergies between biology and materials science will only arise from a deep mutual understanding between disciplines. It will not only be important to optimise the bacterial agent used for a particular application, but also to adjust and enhance the composition of the material to be modified via MICP to be conducive to microbial activity while maintaining its original properties.

## Methods

### Cultivation of bacteria

*E. coli* and *B. subtilis* strains were routinely grown in Lysogeny Broth (LB) at 37°C, 200 rpm and with respective antibiotics. *B. paralicheniformis* and *S. pasteurii* were grown at 30°C, 150 rpm on LB medium and LB medium plus 20g/l urea, respectively. Recombinant *Bacillus subtilis* strains were grown in LB at 30°C, 150 rpm and with respective antibiotics. Optical densities of cultures were routinely measured in a Novaspec Pro spectrometer (Biochrom Ltd, United Kingdom) at 600 nm wavelength in cuvettes of 1 cm light path length (OD_600_). For *B. subtilis* strains with recombinant urease genes, glycerol stocks were prepared by scraping off cells from growth on solid media and re-suspending in autoclaved dH_2_O before adding glycerol to a final concentration of 20 % (w/v). Biofilm promoting medium LBGM contained LB supplemented with 1 % (w/v) glycerol and 0.1 mM MnSO_4_ (36). Where antibiotic selection was required, additions of 10 µg/ml kanamycin, 5 µg/ml chloramphenicol, 100 µg/ml spectinomycin, 20 µg/ml Zeocin or MLS (1 µg/ml erythromycin + 25 µg/ml linocomycin) for *Bacillus subtilis*, or 100 µg/ml ampicillin for *E. coli* were made. For strains requiring induction of heterologous gene expression, media were supplemented with 0.2 % (w/v) xylose.

### Strain construction

All bacterial strains used in this study are listed in Table 1. Plasmids used in this study are listed in Table 2, including details of their construction; primers are listed in Table 3.

**Table 1.**
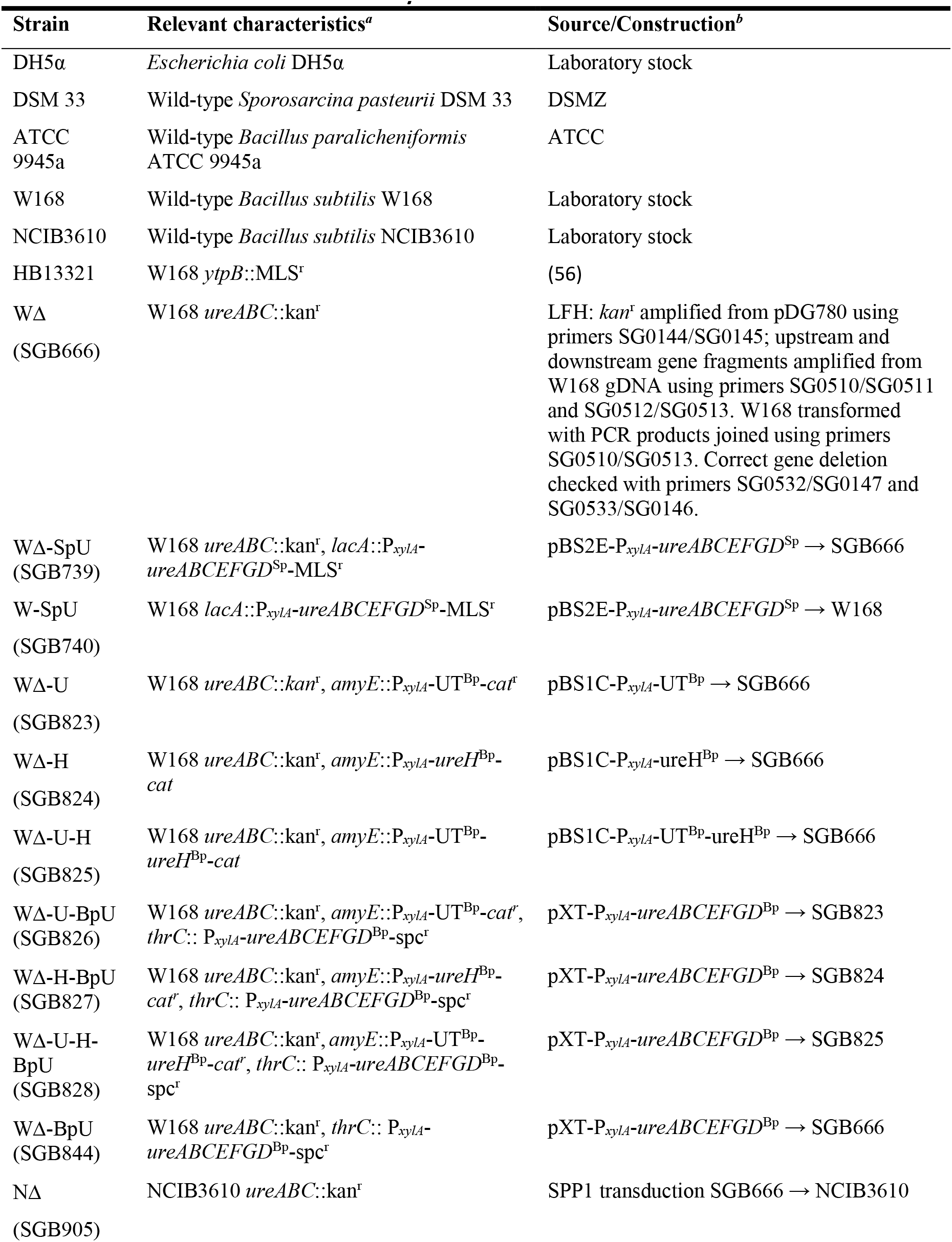
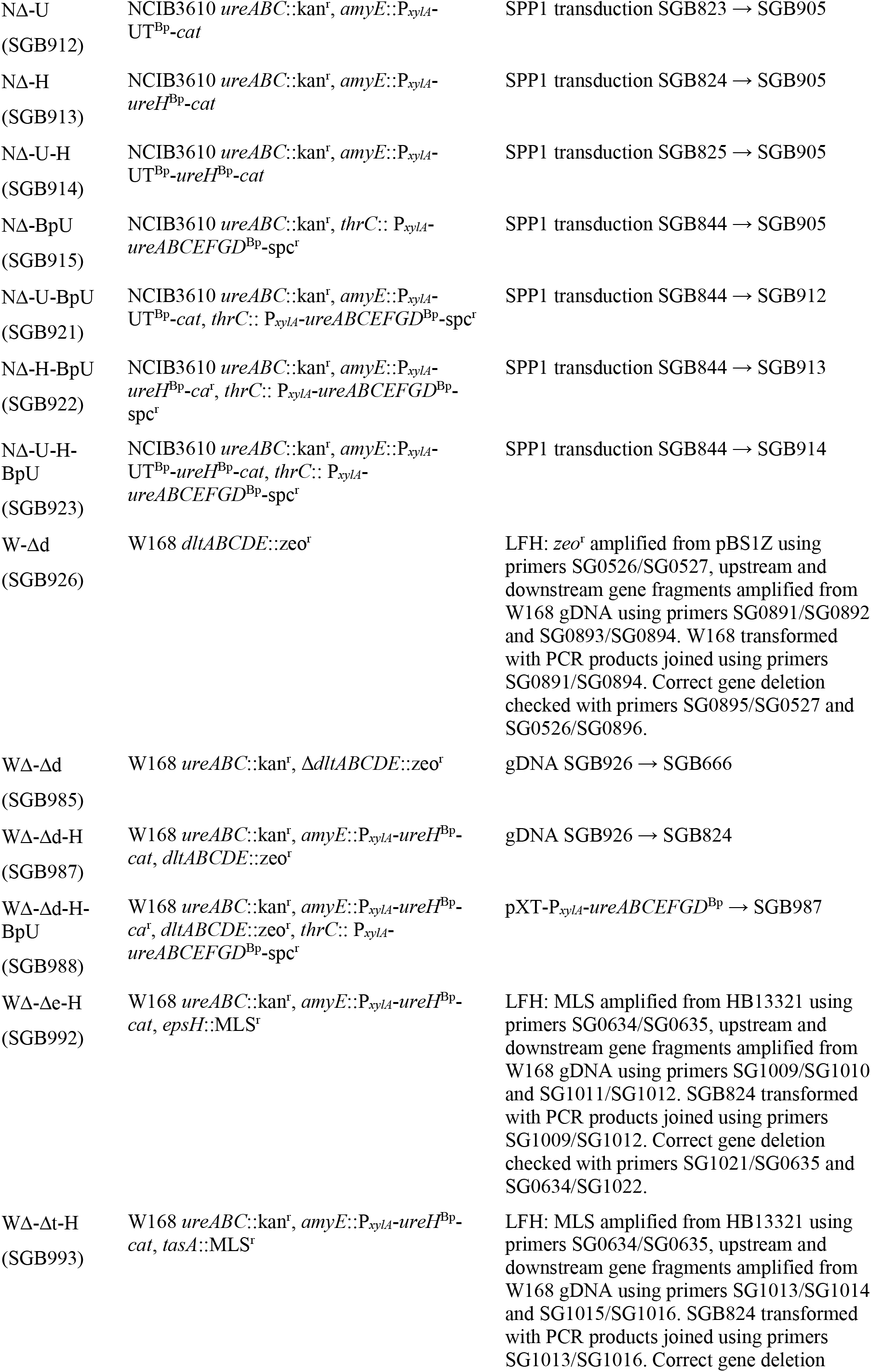
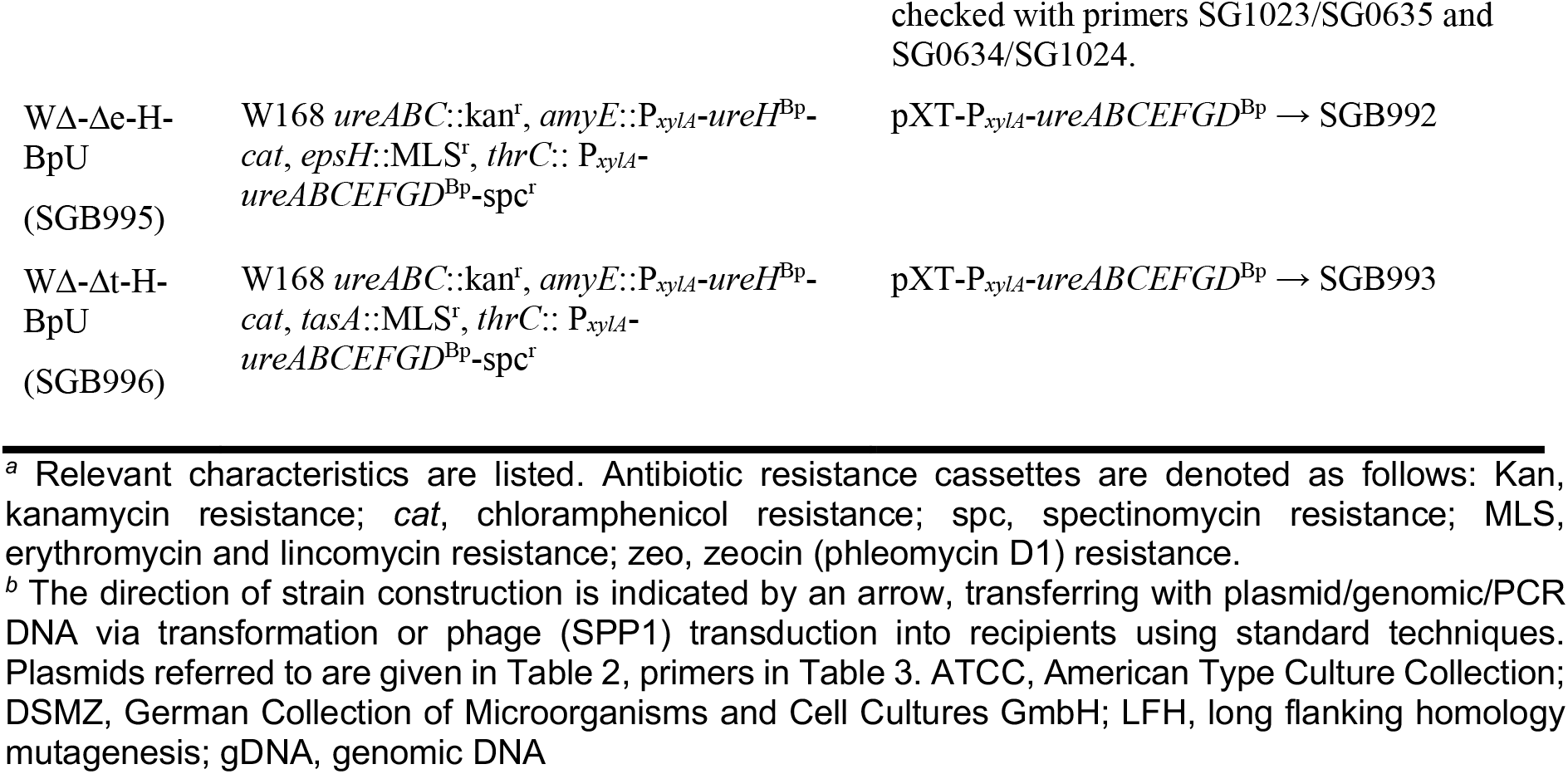
Bacterial strains used in this study.

**Table 1.**
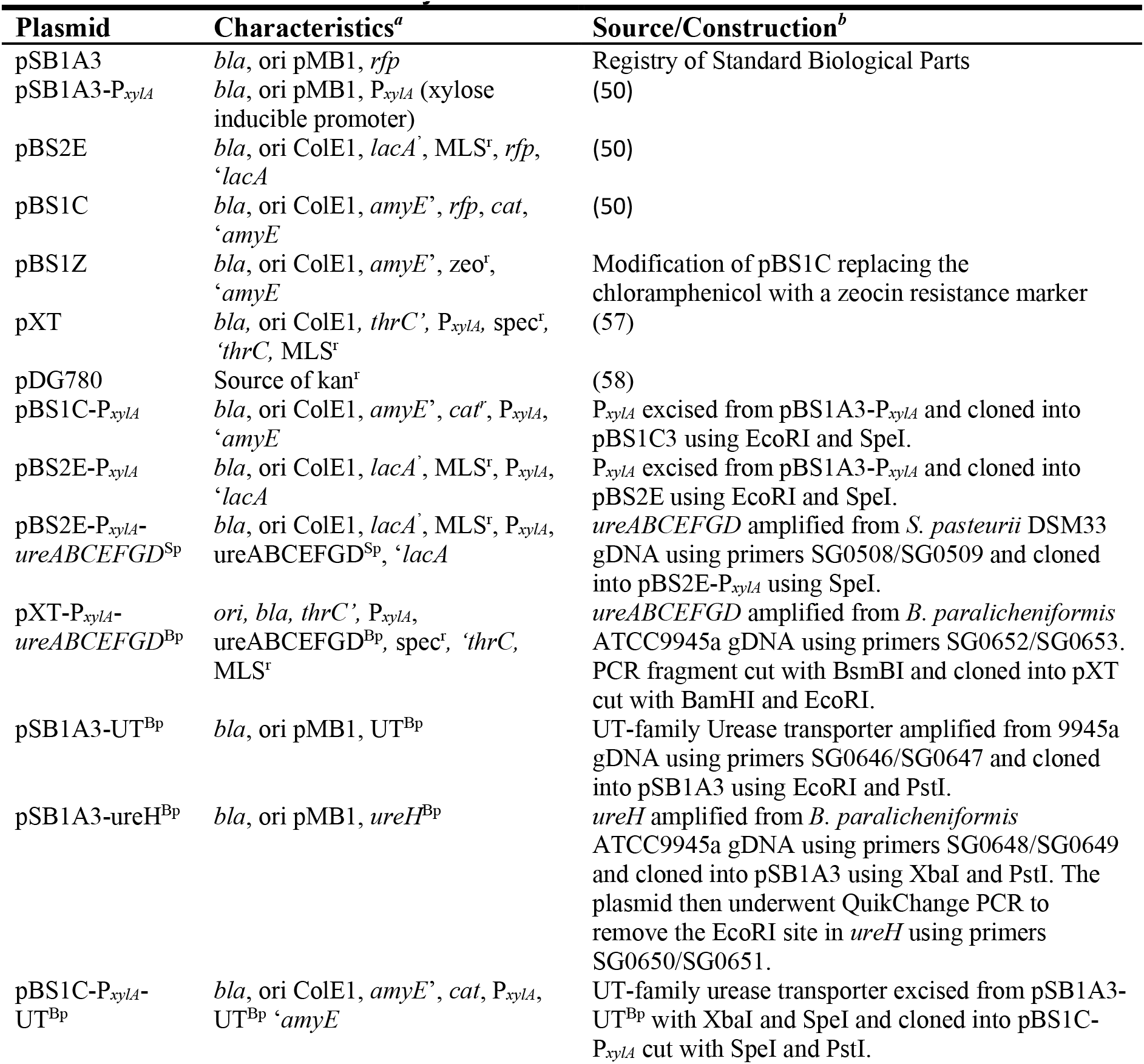
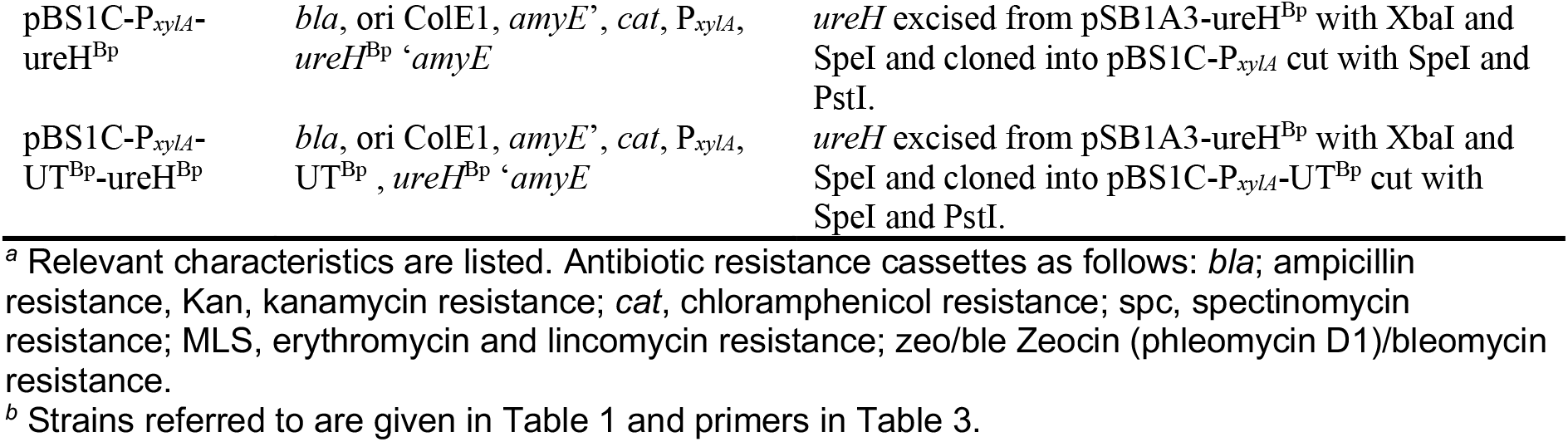
Plasmids used in this study.

**Table 3.**
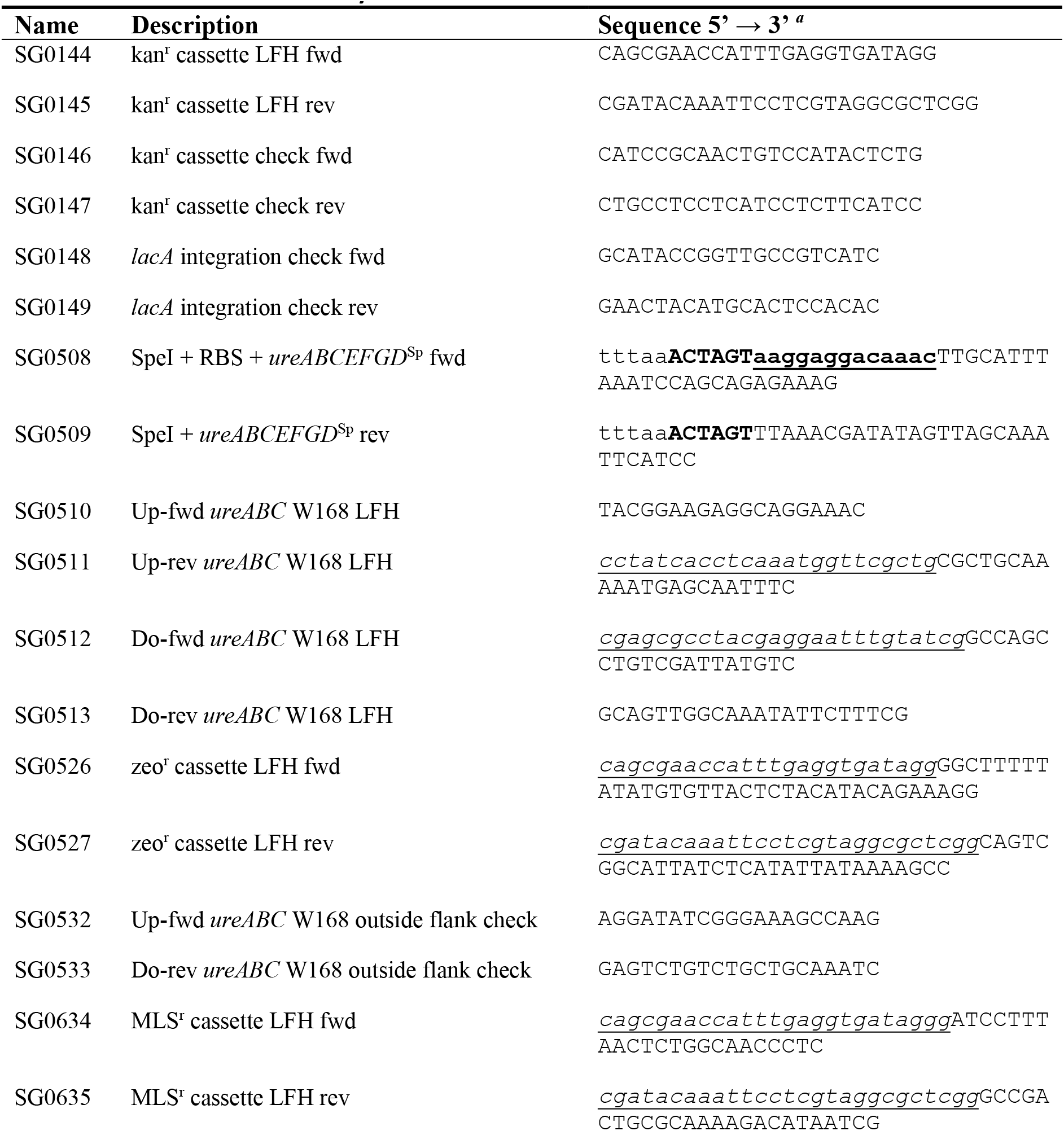
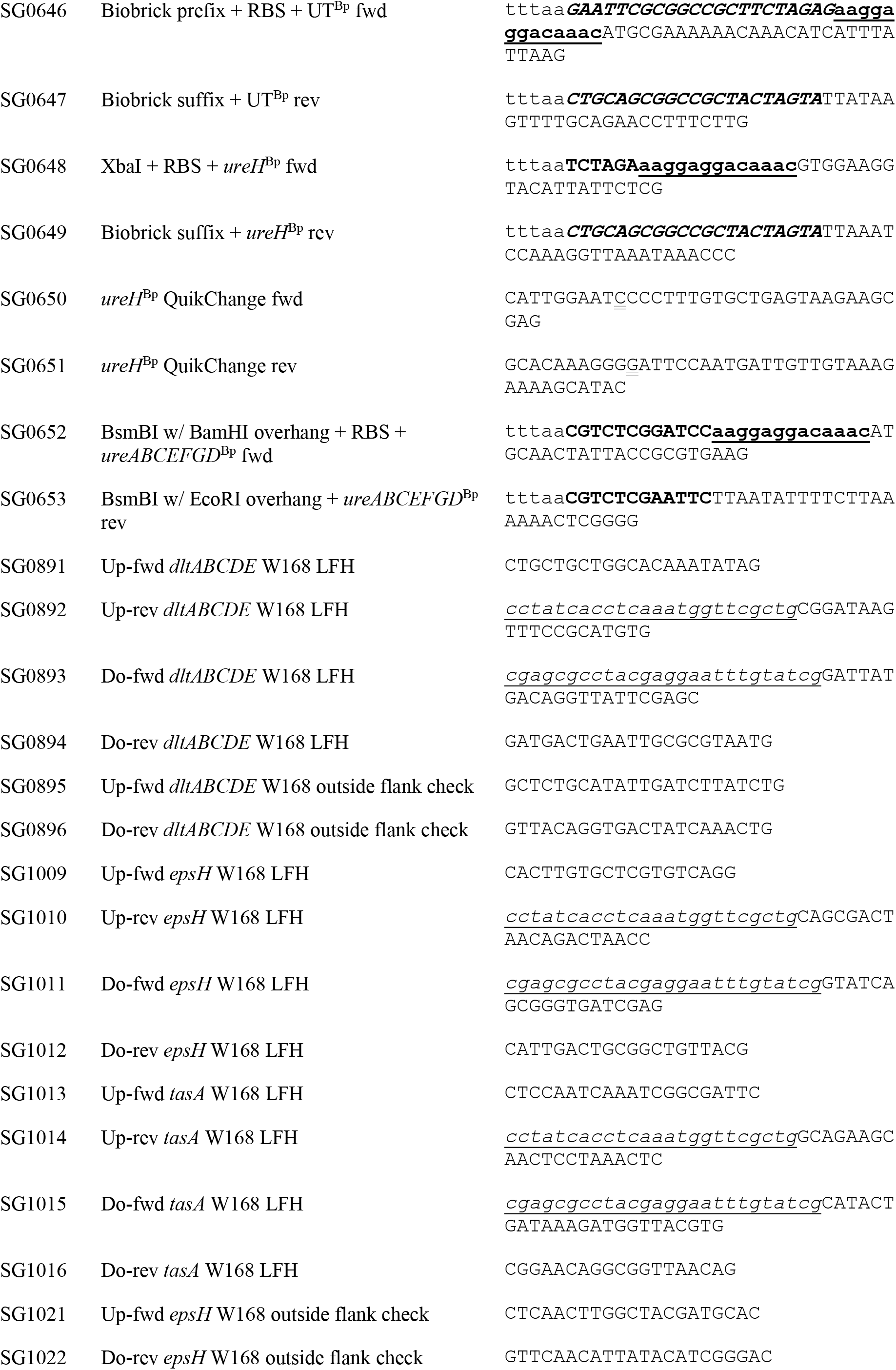
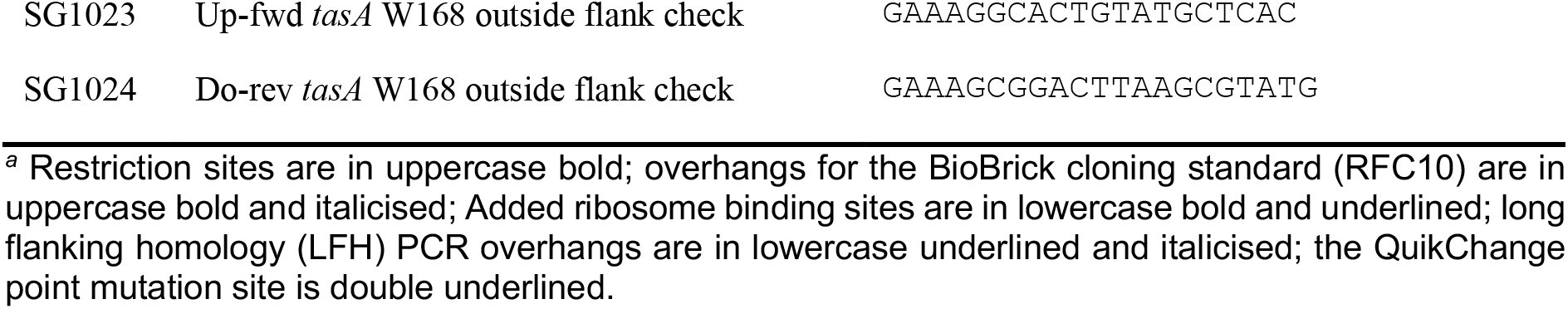
Primers used in this study.

Transformation of *B. subtilis* W168 with plasmid DNA, PCR products or genomic DNA followed induction of natural competence by growth in MNGE medium as previously described (49). Double-crossover integration of the DNA into *B. subtilis* W168 genome at the *amyE* locus was checked by loss of the ability to degrade starch in solid media as described (50). Double-crossover integration of the DNA at the *lacA* locus was checked via colony PCR using primers SG0148/SG0149. Double-crossover integration into the *thrC* locus was checked as threonine auxotrophy during growth in MNGE media with and without addition of 5 mg/ml threonine.

*B. subtilis* W168 gene deletions were constructed by replacing the gene of interest with an antibiotic resistance cassette using long flanking homology mutagenesis as previously described (51).

Successful disruption of the gene was checked via colony PCR using respective primers for outside of the upstream and downstream homology regions as indicated in Tables 1 and 3.

*B. subtilis* NCIB3610 derived strains were constructed by SPP1 mediate phage transduction from *B. subtilis* W168 strains harbouring the mutation/insertion of interest as previously described by Kearns and Losick without DNase treatment of the supernatant (52).

### Zeta potential measurements

The protocol was adapted from Soon *et al.* (53). In brief, cells were sub-cultured in 10 ml LB inoculated to OD_600_=0.05 from an overnight culture and grown at 37°C with agitation until OD_600_=0.5. Cells from 4 ml culture were pelleted at 4,000 ×*g* for 5 min, washed once in 1 volume autoclaved dH_2_O, re-suspended in autoclaved dH_2_O and adjusted to a final OD_600_=0.5. The zeta potential of 1 ml cell suspension was measured in DTS1061 cuvettes with a Zetasizer Nano ZS equipped with a 633 nm red laser (Malvern Instruments Ltd, Malvern, UK) using the Helmholtz-Smoluchowski theory. Measurements were performed at 25°C (120s calibration) and taken as triplicate readings (technical replicates) and repeated for three biological replicates.

### Nisin-dependent growth inhibition

This assay was adapted from concentration-dependent killing experiments (54). Overnight cultures grown at 37°C were adjusted to OD_600_=1 with sterile dH_2_0, and 15 μl were used to inoculate 150 μl of LB in clear flat-bottom 96-well plates. Cells were grown in a TECAN-Spark microplate reader (Tecan Group Ltd., Switzerland) at 37°C and orbital shaking at 180 rpm. In late exponential phase (OD_600_=0.5, corresponding to OD_600_=1.37 in a 1 cm standard cuvettes), nisin was added to final concentrations of 0-64 μg/ml, followed by continued monitoring of OD_600_ every 15 min over 5 hours. Killing by nisin was assessed as a decrease in OD_600_ over time relative to the control without antibiotic addition.

### Cytochrome C binding assay

The cytochrome C binding assay was adapted from Revilla-Guarinos *et al.* (39). Overnight cultures (approximately 2 ml of OD_600_=2.5) were harvested by centrifugation (2 min at 16,000 ×*g*) and re-suspended in 1.8 ml 20 mM MOPS-NaOH pH 7 and adjusted to an OD_600_=2.5. Cells were suspended by thorough mixing and then incubated with a final concentration of 250 μg/ml cytochrome C for 10 min at room temperature. Cells were removed by centrifugation (4 min at 16,000 ×*g*) and the concentration of unbound cytochrome c remaining in the supernatant was measured as absorbance at 530 nm wavelength (A_530_) in a spectrophotometer. This value was compared to the A_530_ value of a pure 250 µg/ml cytochrome C solution without bacterial cell (the 0 % binding bench mark) to calculate the binding percentage.

### Calcite precipitation assays

Calcite precipitation on bacterial colonies was assessed on LBC, LBGMC, and B4 solid precipitation media. LBC and LBGMC media were LB or LBGM, respectively, supplemented with 10 g/L calcium acetate. B4 precipitation media contained 0.4 g/L yeast extract, 1 g/L fructose, 0.25 g/L calcium acetate, adjusted to pH 8 with NaOH, adapted from the original composition by changing the sugar source (31). For ureolytic strains, precipitation media were supplemented with 20 g/L urea. To prepare the inoculum, cells were grown overnight on solid LB medium with antibiotics at 30°C and then scraped from the agar and resuspended to OD_600_=0.5 in autoclaved dH_2_O. Of this, 10 µl was spotted onto agar plates of either LBC, LBGMC or B4 media. Plates were incubated at 30°C for 1-2 weeks, imaging the colony and crystal formation at regular intervals at 5× magnification with a Leica MZ7.5 stereomicroscope (Leica, United Kingdom), using an Infinity 2 Lumenera microscope camera (Teledyne Technologies Inc., United States).

### Electron microscopy

To examine crystal morphology and composition of the mineral precipitate, crystals were carefully scraped from the bacterial colonies that underwent the calcite precipitation assay. Crystals were repeatedly washed in volumes of 1 ml dH_2_O by gentle pipetting to remove any remaining cells and agar until the water remained clear and then dried overnight at room temperature to evaporate any remaining liquid. Before imaging, samples were placed under vacuum overnight. For Scanning Electron Microscopy (SEM), samples were coated with gold for 1-3 min using an Edwards S150B sputter coater (Edwards Group Ltd., United Kingdom) and imaged on a JOEL JSM-6480LV SEM microscope (JEOL Ltd., Japan). An energy dispersive x-ray analyser (EDX) equipped on the SEM was used for elemental analysis of the calcite crystals prior to gold coating.

### Image manipulation

Photographs presented in this study were taken with either a hand-held camera, microscope-mounted cameras or generated by the SEM software and were only edited for clarity of display, i.e. cropping of excess image area and correction of brightness and contrast.

### Urease activity assays

To determine urease activity, rapid urease test broth (RUB) containing 0.1 g/l yeast extract, 20 g/l urea, 0.67 mM monopotassium phosphate, 0.67 mM disodium phosphate, 0.01 g/l phenol red, pH 6.8±0.2, was used. For qualitative urease assessment, 2 ml RUB was inoculated with colonies picked directly from solid growth media. Suspensions were subsequently incubated at 37°C, 200 rpm. The semi-quantitative urease assay was based on a protocol by Okyay and Rodrigues (55). Cells were grown overnight on their respective standard solid media with antibiotics at 30°C, scraped off the agar and suspended in autoclaved dH_2_O to OD_600_=0.5. Of this, 20 µl were used to inoculate 200 µl of RUB in clear flat-bottom 96-well plates, followed by incubation in a TECAN-Spark microplate reader (Tecan Group Ltd., Switzerland) at 30°C and orbital shaking 180 rpm. Absorbance readings were taken at 560 nm wavelength (A_560_) every 15 min for 24 hours to assess the change in pink colour formation over time as a proxy for whole cell enzyme activity. To ascertain that no cell growth occurred that could have affected the A_560_ readings, OD_600_ was monitored in the same way. As there were no changes in OD_600_ over time, wells containing RUB without cells were used as a blank for the A_560_ readings.

To assess urease activity on solid media, 0.01 g/l phenol red were added, and the media were adjusted to an initial pH of 7.2.

## List of Abbreviations

EDX: Energy-dispersive X-ray spectroscopy
LBC: LB-Calcium medium
LBGM(C): LB-Glycerol-Manganese (-Calcium) medium
MICP: Microbially induced calcite precipitation
RUB: Rapid urea test broth
SEM: Scanning Electron Microscopy

## Funding

This work was funded through the Engineering and Physical Sciences Research Council (EPSRC; EP/PO2081X/1) Resilient Materials for Life (RM4L) project with support from industrial collaborators/partners. TDH was supported by a University of Bath Research Studentship. The funding bodies did not have any role in design of the study or collection, analysis, and interpretation of data or in writing the manuscript.

## Authors’ contributions

TDH planned and carried out all experimental work, analysed and interpreted the data, prepared the figures and co-wrote the manuscript. SG supervised the work, contributed to study design and data interpretation and co-wrote the manuscript. KP provided materials science insights, co-supervised the work and edited the manuscript. All authors read and approved the final manuscript.

## Acknowledgements

The authors gratefully acknowledge the Technical staff within the Department of Biology and Biochemistry as well as the Material and Chemical Characterisation team at the University of Bath for technical support and assistance in this work

---

**Additional file 1.**
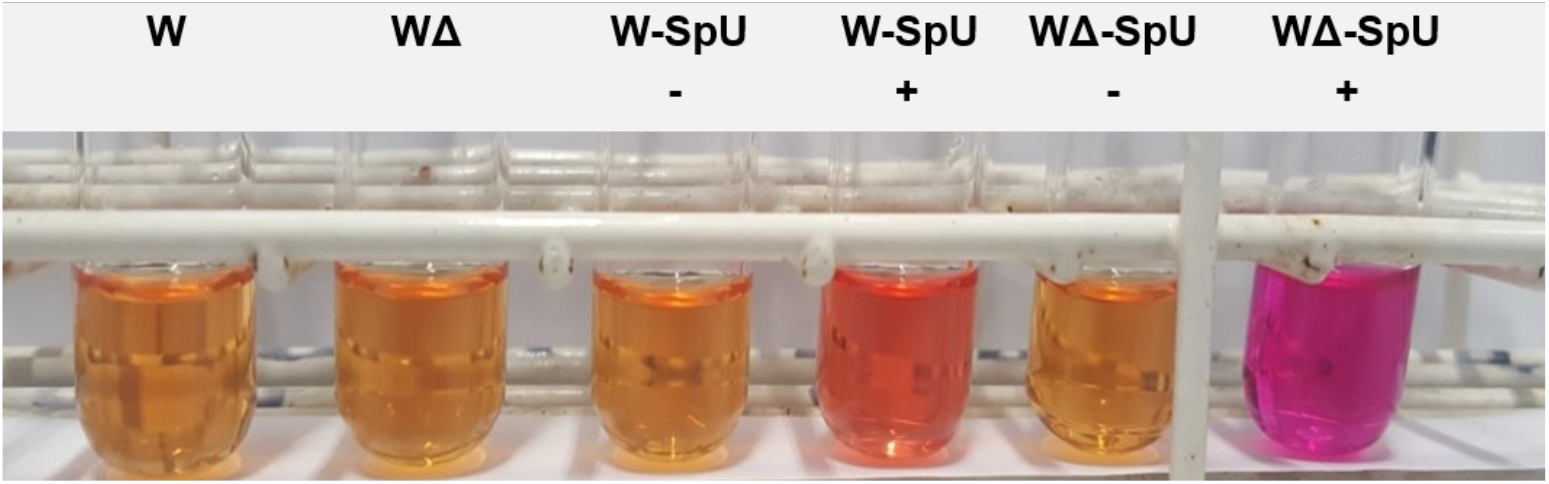
Heterologous expression of *Sporosarcina pasteurii* urease in *Bacillus subtilis.* Strains of *B. subtilis* W168 (W) or a derived *ureABC-deletion* strain (WΔ) were transformed with a xylose-inducible plasmid containing the *S. pasteurii* urease gene cluster (SpU). Rapid urease test broth with or without addition of 0.2% xylose (+, -) to induce urease gene expression were inoculated with cells taken from growth on solid media and incubated at 37°C for 2 days. Urease activity is observed as a yellow-to-pink colour change of the medium resulting from a pH increase due to the release of ammonia from urea.

**Additional file 2.**
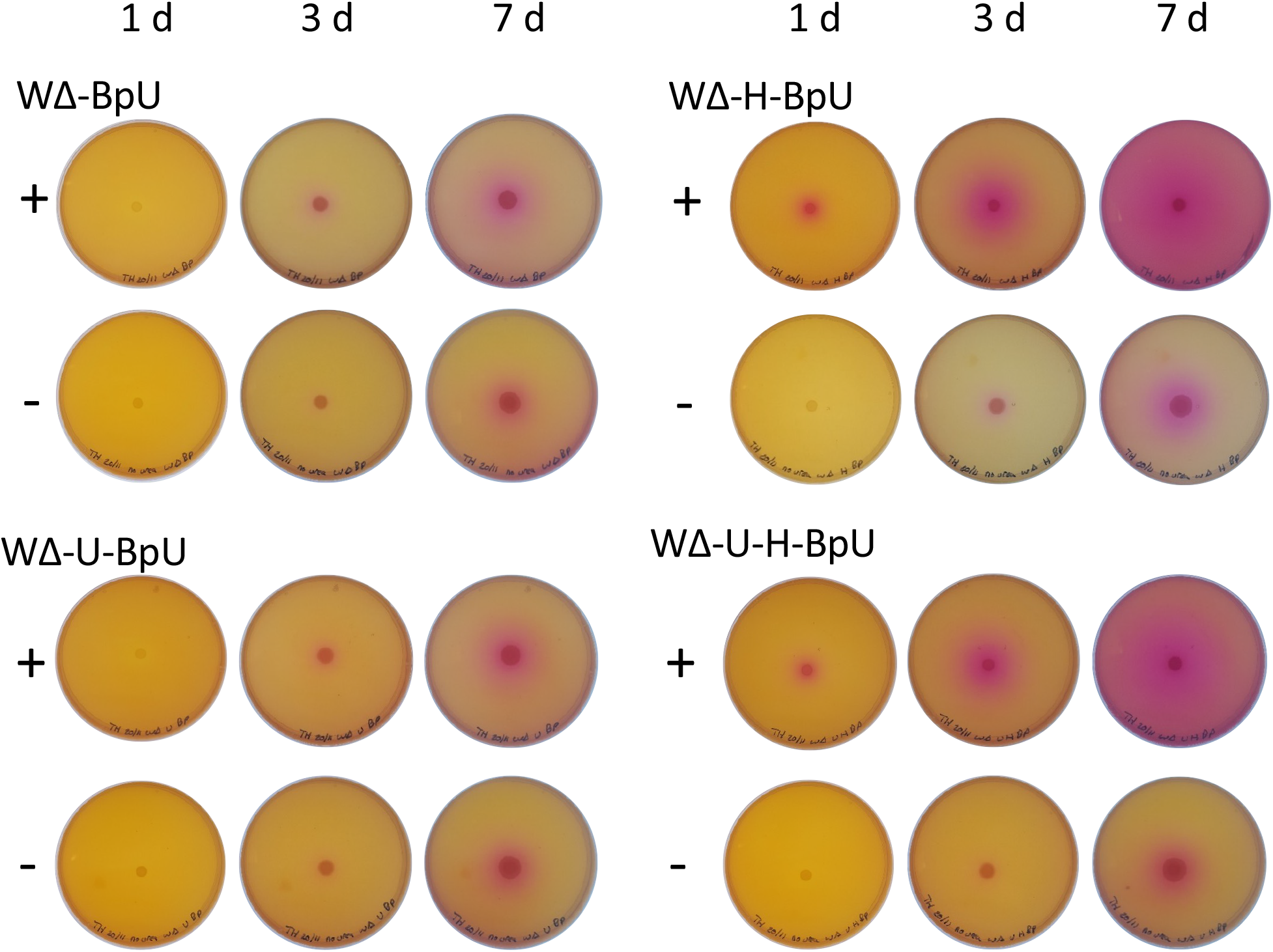
Urease activity on solid media following heterologous expression of *Bacillus paralicheniformis* urease genes in *B. subtilis*. Cells of a *ureABC* deletion strain of *B. subtilis W168* (WΔ) containing expression constructs for *B. paralicheniformis ureABCEFGD* (BpU), *ureH* (H), UT (U) or combinations thereof were spotted onto LBC agar plates with (+) or without (-) urea. The plates contained phenol red and had been adjusted to an initial pH of 7.2. Plates were incubated at 30°C and photographed after 1, 3 and 7 days as indicated. Urease activity is visible as a stronger development of pink colouration as compared to the urea-free control; the slight pink colouration immediately surrounding the bacterial growth is due to general metabolic activity. Results shown are representative of three independent experiments.

**Additional file 3.**
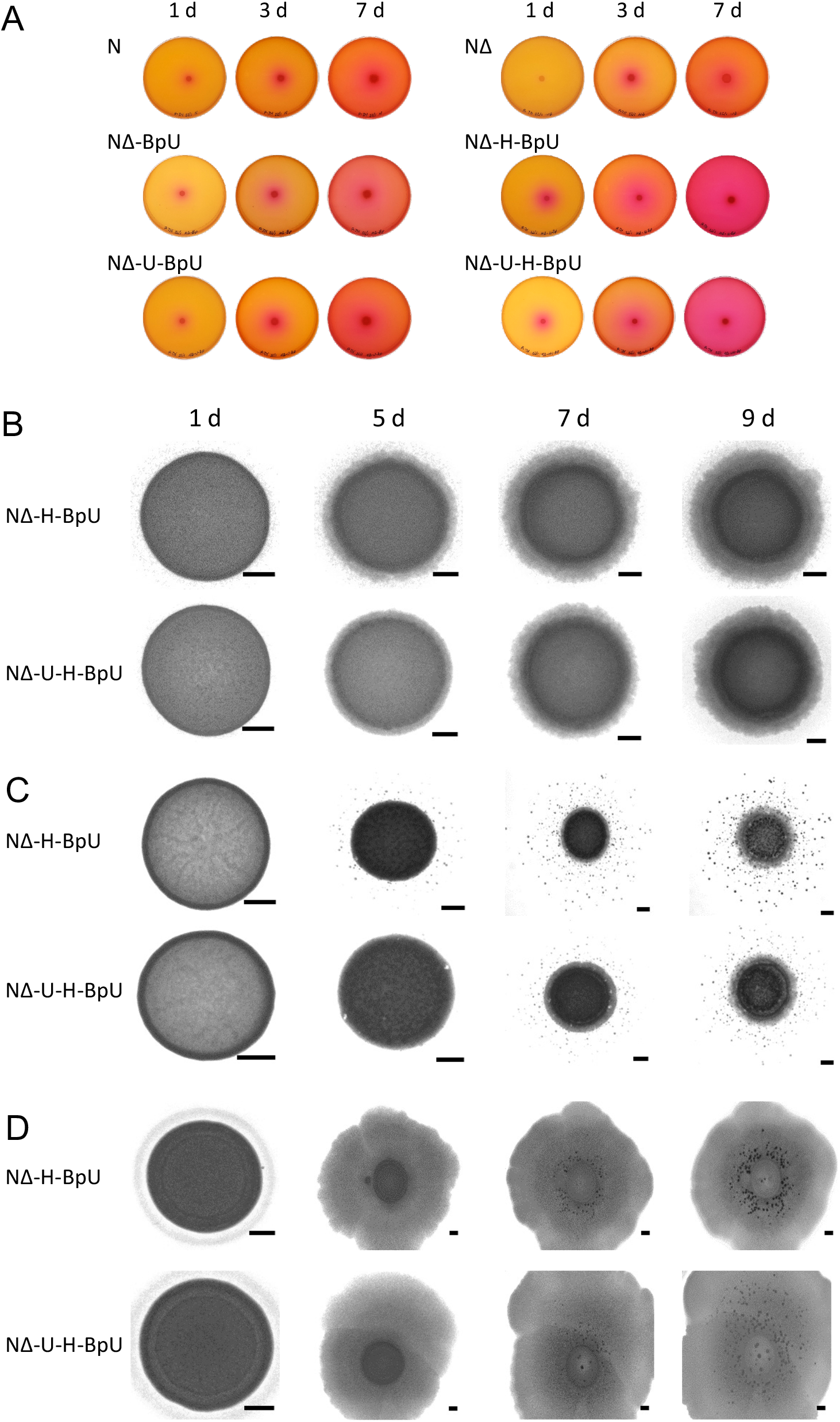
Engineered urease and biomineralisation activity in *B. subtilis* NCIB3610. **(A)** urease activity assay on LBC medium containing urea, xylose, phenol red and adjusted to an initial pH of 7.2. Cells were spotted onto the centre of the plate and photographed over seven days of incubation at 30°C as indicated above. Urease activity was observed as pink colouration of the agar. **(B-D)** Biomineralisation assays on LBC medium (B), B4 medium (C) or LBGMC biofilm-promoting medium (D), all supplemented with urea and xylose. Cells were spotted onto the centre of the agar, incubated at 30°C for up to 9 days and imaged with a stereomicroscope at the time points indicated above in panel B. The image area shown varies between days to allow visualisation of mineral crystals outside the colony area in panel C or to accommodate the increased area of growth in panel D. Scale bars represent 1 mm in size. Representative results of two independent repeats are shown. In all panels, the strain nomenclature is as follows: wild-type *B. subtilis* NCIB3610 (N); its isogentic *ureABC* deletion strain (NΔ); derived strains containing expression constructs for *B. paralicheniformis ureABCEFGD* (BpU), *ureH* (H), UT (U) or combinations thereof.

**Additional file 4.**
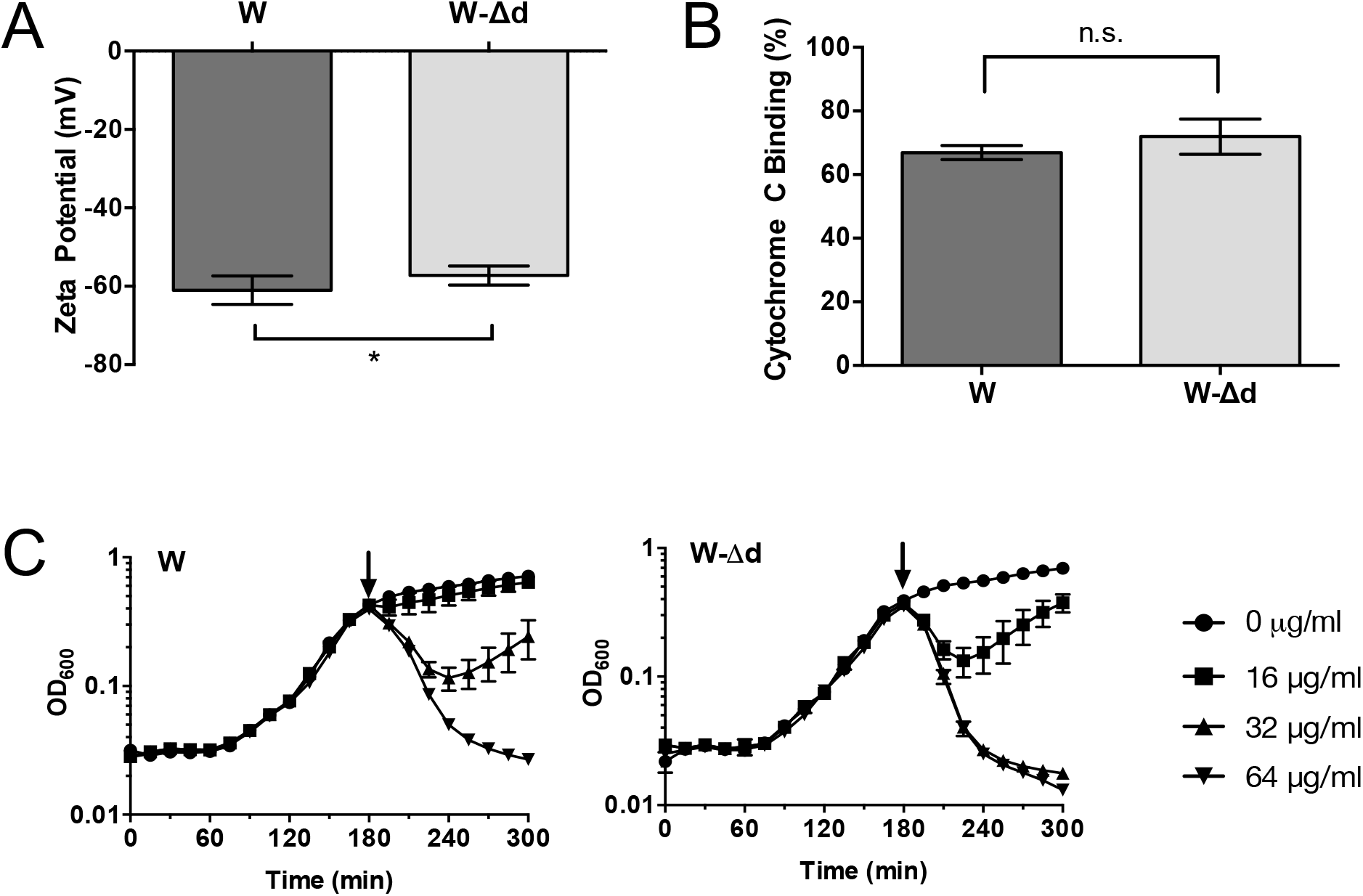
Effects of *dtlABCDE* deletion on the cell surface charge of *B. subtilis*. Cells of wild-type *B. subtilis* W168 (W) or an isogenic *dltABCDE* deletion strain were subjected to surface charge analyses. **(A)** Zeta potential measurements. Cells were grown to exponential phase (OD_600_ = 0.5-1), harvested, washed and resuspended in dH_2_O to OD_600_ = 0.5, and the zeta potential measured on a Zetasizer Nano ZS at 25 °C. Data are shown as mean ± standard deviation of three biological repeats, each measured in technical triplicates. n.s. shows p>0.05 and * shows p≤0.05 from an un-paired t-test analysis. **(B)** Cytochrome C binding assay. Overnight cultures were resuspended in MOPS/NaOH [pH7] to OD_600_ = 2.5 and incubated with 250 µg/ml cytochrome C for 10 minutes at room temperature. Percentage binding was calculated as absorbance difference of the supernatant relative to samples without bacteria. Data are shown as mean ± standard deviation of three to four biological repeats; n.s. p>0.05 in an unpaired t-test analysis. **(C)** Nisin-dependent killing assay. Cells were grown in LB medium in a Tecan microplate reader to OD_600_ = 0.4-0.5. At the time point shown by the arrows, nisin was added at the indicated concentrations and OD_600_ monitored over time. Data are shown as mean and standard deviation of three technical repeats and are representative of three biological repeats.

## Notes

### Competing Interest Statement

The authors have declared no competing interest.

